# Engagement of sialylated glycans with Siglec receptors on myeloid suppressor cells inhibit anti-cancer immunity via CCL2

**DOI:** 10.1101/2023.06.29.547025

**Authors:** Ronja Wieboldt, Emanuele Carlini, Chia-wei Lin, Anastasiya Börsch, Andreas Zingg, Didier Lardinois, Petra Herzig, Leyla Don, Alfred Zippelius, Heinz Läubli, Natalia Rodrigues Mantuano

## Abstract

Overexpression of sialic acids on glycans, called hypersialylation is a common alteration found in cancer. Hypersialylation can, for example, enhance immune evasion via interaction with sialic acid-binding immunoglobulin-like lectin (Siglec) receptors on tumor-infiltrating immune cells. Here, we tested the role of sialic acid on myeloid-derived suppressor cells (MDSCs) and their interaction with Siglec receptors. We found that MDSCs derived from the blood of lung cancer patients and tumor-bearing mice strongly express inhibitory Siglec receptors. In murine cancer models of emergency myelopoiesis, Siglec-E knockout on myeloid cells resulted in prolonged survival and increased infiltration of activated T cells. Targeting suppressive myeloid cells by blocking Siglec receptors or desialylation led to strong reduction of their suppressive potential. We further identified CCL2 as mediator involved in T cell suppression upon interaction of sialoglycans and Siglec receptors on MDSCs. Our results provide mechanistic insights how sialylated glycans inhibit anti-cancer immunity by facilitating CCL2 expression.

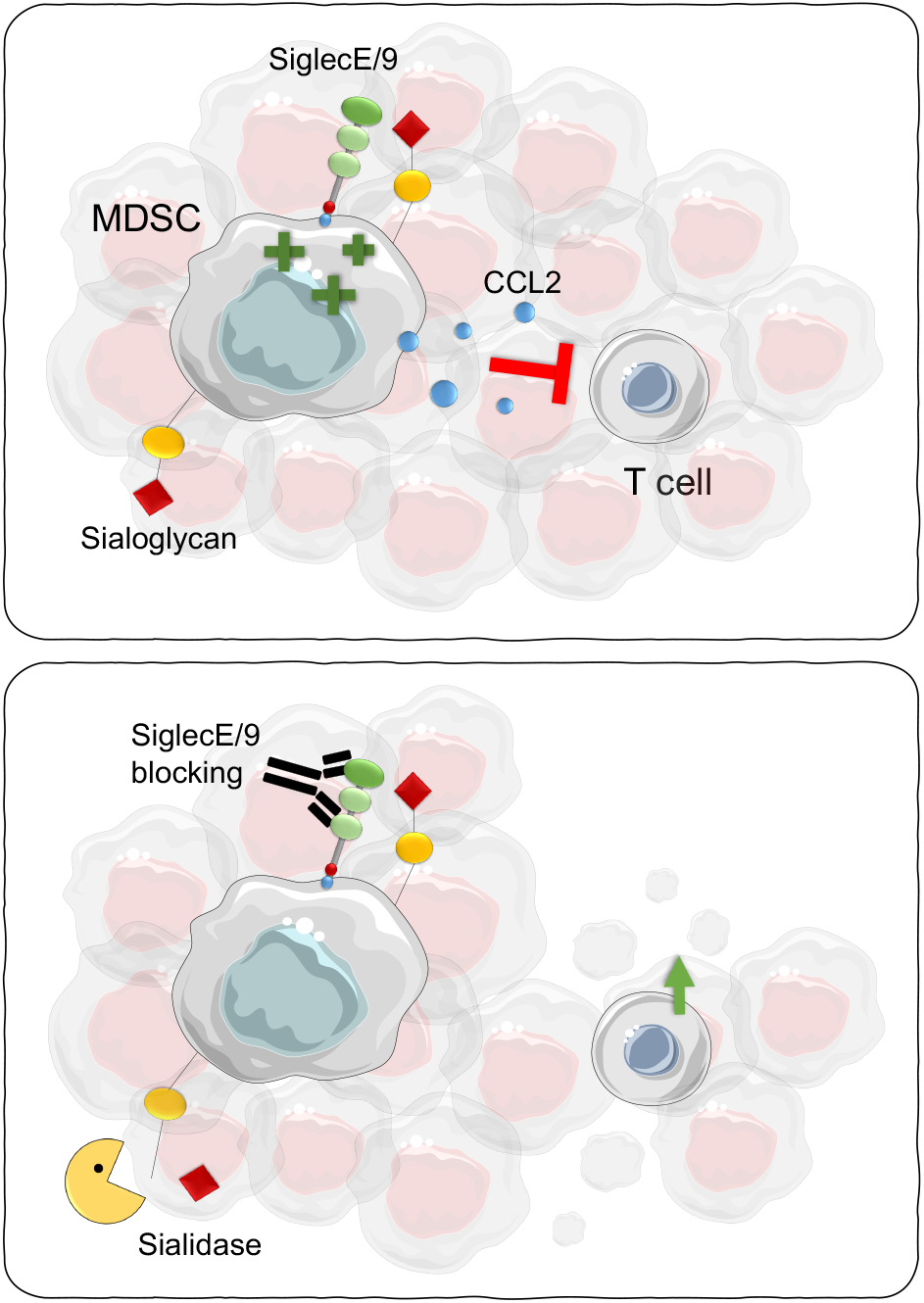

## Introduction

Immunotherapy has revolutionized the field of cancer treatment, but only a subset of patients experiences an anti-tumor response and many relapse over time. Primary or acquired therapy resistance can be facilitated by a suppressive tumor microenvironment (TME) helping cancer cells to evade immunosurveillance (1). For example, overexpression of sialic acid-containing glycans (sialoglycans) lead to hypersialylation. Hypersialylation can drive cancer progression by mediating stability of cell surface receptors of cancer cells, integrin-mediated interactions and enhance immune invasion by engaging Sialic acid-binding immunoglobulin-like lectin (Siglec) receptors on immune cells, modulating antigen presentation, interacting with selectin receptors and facilitating stability of surface proteins on immune cells (2–4). Siglec receptors are highly expressed by different immune cells. The majority of Siglec receptors are part of the CD33-related inhibitory Siglec family which mainly signal via cytosolic immunoreceptor tyrosine-based inhibitory motifs (ITIMs) (5). Recent studies highlight the importance of inhibitory Siglecs for the regulation of various immune cells including T cells, natural killer (NK) cells, dendritic cells (DCs) and macrophages (6–9). Targeting the Siglec-sialoglycan axis in cancer presents a potential immune checkpoint for cancer immunotherapy that can be achieved by Siglec blocking or cleaving of sialoglycans using sialidase (10). For example, a sialidase linked to a tumor-directed antibody was successfully used *in vivo* and led to better tumor control by repolarization of macrophages in the TME (11,12).

Pathologically activated myeloid cells are known to promote an immunosuppressive TME and mark a promising target for cancer therapy (13). This closely-related family of myeloid suppressors mainly consist of tumor-associated macrophages (TAM) and myeloid derived suppressor cells (MDSCs). Induced by aberrant myelopoiesis, MDSCs are a heterogenous group of immature myeloid cells which are involved in suppression of various immune cells via secretion of suppressive cytokines or direct cell-cell interactions (14). MDSCs can be subdivided in monocytic MDSCs (mMDSCs) and granulocytic or polymorphonuclear MDSCs (gMDSCs or PMN-MDSCs), which are identified according to their monocytic or granulocytic myeloid cell lineage markers (15). Apart from Siglec-3/CD33 as phenotypic marker expressed by all MDSCs, recent studies further describe the expression of Siglec-5, Siglec-7 and Siglec-9 on human glioma patient MDSCs and Siglec-E on murine MDSCs (11,16). Nevertheless, little is known about the purpose and function of Siglecs and sialoglycan ligands on MDSCs and their expression in health and disease. Here, we investigate the role of the Siglec-sialoglycan axis for MDSC generation and function in the TME in humans and mice and their expression patterns in lung tumor patients and healthy individuals.

## Results

### 1. Myeloid cells express inhibitory CD33-related Siglecs in cancer

Although no exclusive MDSC markers are known, MDSCs can be identified by co-expression of phenotypic markers which vary between murine and human. All murine MDSCs express Gr1 and CD11b and can be subdivided in Ly6G^+^Ly6C^low^ gMDSCs and Ly6C^high^ mMDSCs (17). In humans, gMDSCs are phenotypically characterized as CD33^+^CD11b^+^HLA-DR^low/-^CD15^+^CD14^-^ and mMDSCs as CD33^+^CD11b^+^HLA-DR^low/-^ CD15^-^ CD14^+^ (15). All human MDSCs express Siglec-3/CD33 as a phenotyping marker (15), but little is known about the expression of additional Siglec receptors on MDSCs in human cancer patients and their functional relevance (16).

Here, we investigate the expression of Siglec receptors found on lung patient-derived MDSCs as well as on myeloid cells from healthy donors identified as lineage^-^ CD33^+^CD11b^+^HLA-DR^low/-^ cells (Supp. 1 A, C). Human MDSCs from lung cancer patients expressed high levels of Siglec-5, Siglec-7, Siglec-9 and Siglec-10 in the periphery and within the tumor (1 A, Supp. 1 C). Increased levels of Siglec-9 and Siglec-10 were expressed on patient MDSCs in the periphery compared to myeloid cells from healthy donors (1 B, C). Siglec-5 and Siglec-7 showed similar expression patterns on MDSCs derived from cancer patients and myeloid cells from healthy controls in peripheral blood (Supp.1 D, E).

**Figure 1:**
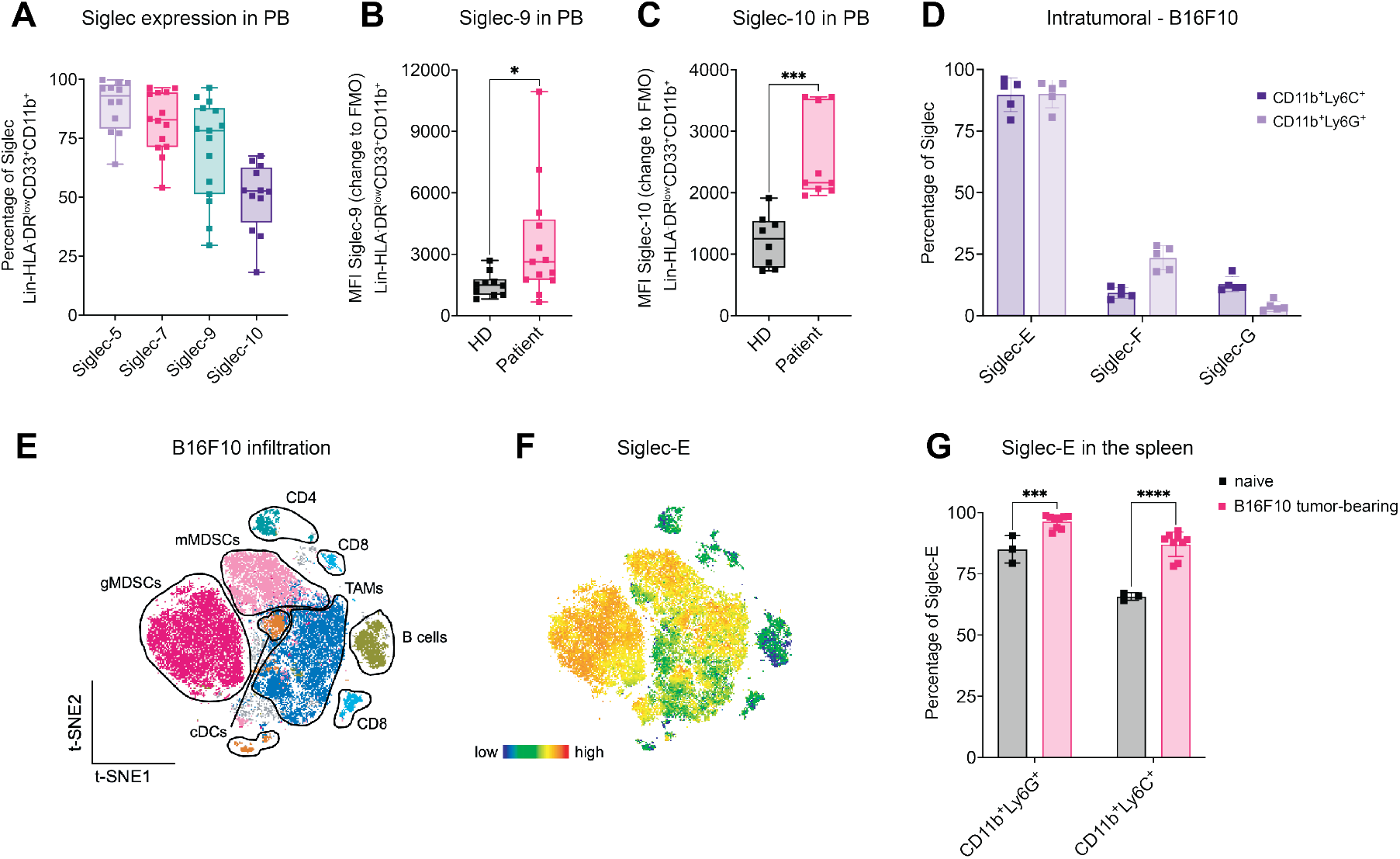
Myeloid cells express Siglecs in human and mice. **(A)** Percentage of Siglec-5, Siglec-7, Siglec-9 and Siglec-10 expressed on Lin^-^ HLADR^low^CD33^+^CD11b^+^ cells detected in peripheral blood (PB) from lung cancer patients by flow cytometry. *N=12-14 donors from N=4 experiment*. **(B)** MFI of Siglec-9 and **(C)** Siglec-10 on CD45^+^Lin^-^HLADR^low^CD33^+^CD11b^+^ cells derived from healthy donor and lung cancer patient PB from (A). MFI is shown as change to FMO and was determined by flow cytometry. *N=8-13 donors with at least N=2 experiments.* **(D)** Subcutaneously injected endpoint tumors from B16F10 melanoma engrafted mice were harvested, digested and immune cell infiltration was assessed by multi-color flow cytometry. Siglec-E, Siglec-F and Siglec-G expression was assessed on CD45^+^CD11b^+^Ly6G^+^ and CD45^+^CD11b^+^Ly6C cells. *N=5 mice.* **(E)** T-distributed stochastic neighbor embedding (t-SNE) projection of multicolor flow cytometry immunophenotyping of pooled infiltrating immune cells from B16F10 tumors. *N=5 mice.* **(F)** Siglec-E expression intensity is shown as color gradient from blue (low) to red (high). **(G)** Spleens from naïve and B16F10 melanoma tumor-bearing mice at endpoint were collected and analyzed for Siglec-E expression via flow cytometry. *N=3-9 mice per group*. Data are presented as mean. Error bar values represent SD. Two-tailed unpaired Student’s t test or multiple unpaired t-tests (G) was used. *P<0.05, **P<0.01, ***P<0.001, and ****P<0.0001.

Next, we investigated whether our findings were similar in mice by analyzing mouse CD33-related inhibitory Siglec receptors on tumor-bearing and naïve MDSCs including Siglec-E, Siglec-F and Siglec-G, which resemble potential functional paralogs of human Siglec-9, Siglec-8 and Siglec-10, respectively (18). To this end, Siglec expression of tumor infiltrating and spleen-derived MDSCs in tumor-bearing and naïve mice was analyzed by assessing CD11b^+^Ly6G^+^ and CD11b^+^Ly6C^+^ populations which in the tumor context are described as gMDSCs and mMDSCs, retrospectively (17) (Supp.1 F). High levels of Siglec-E were identified across infiltrating MDSCs in different tumor types, intermediate levels of Siglec-F were only found on gMDSCs and Siglec-G was rarely expressed on both MDSC types (1 D, Supp.1 G). Siglec-E mean fluorescence intensity (MFI) was the highest expressed on gMDSCs compared to all other tumor-infiltrating immune cells in B16F10, followed by mMDSCs and TAMs (CD11b^+^F4/80^+^) (1 E, F). Furthermore, Siglec-E expression was increased on CD11b^+^Ly6G^+^ and CD11b^+^Ly6C^+^ populations from tumor-bearing mice in the spleen compared to naïve littermates in B16F10 and EL4-bearing mice (1 G, Supp.1 H). These results show an increased expression of inhibitory Siglecs in human and mice in the cancer context.

### 2. Myeloid cells in cancer are hypersialylated

Binding of Siglec receptors to their sialoglycan ligands has been described before and is involved in immune cell interactions with cancer cells but also during antigen presentation and formation of adaptive immunity (19). Although hypersialylation is a common hallmark of cancer and is mainly studied on cancer cells, sialoglycans can additionally be expressed on secreted glycoproteins, glycolipids and the cell surface of immune cells themselves (20,21). To investigate the potential interactions of Siglec receptors with ligands on the surface of MDSCs, we further assessed the sialylation pattern of MDSCs in the TME. Lectin staining was performed to assess surface sialoglycan ligands including Sambucus Nigra Lectin (SNA) detecting α-2,6-linked sialic acids and Maackia Amurensis Lectin II (MALII) detecting α-2,3-linked sialic acids. Both lectins showed a significantly increased staining on peripheral MDSCs derived from lung cancer patients compared to myeloid cells from healthy donors (2 A, B). No changes were detected in Peanut Agglutinin (PNA) levels, a galactosyl (β-1,3) N-acetylgalactosamine structure that is usually masked by sialic acid binding (Supp.2 A). In addition, mass spectrometry(MS)-based analysis on released N-glycans from peripheral patient-derived CD33^+^ cells showed a clear increase of terminally sialylated N-glycans containing multiple sialic acids when compared to myeloid cells from healthy donors (2 C, D). The main N-glycan structure found on healthy donor CD33^+^ cells were core-fucosylated N-glycans with mono sialic acid and PolyLacNAc. In addition, N-glycans containing more than 2 fucoses were also observed, indicating the presence of Lewis structures.

**Figure 2:**
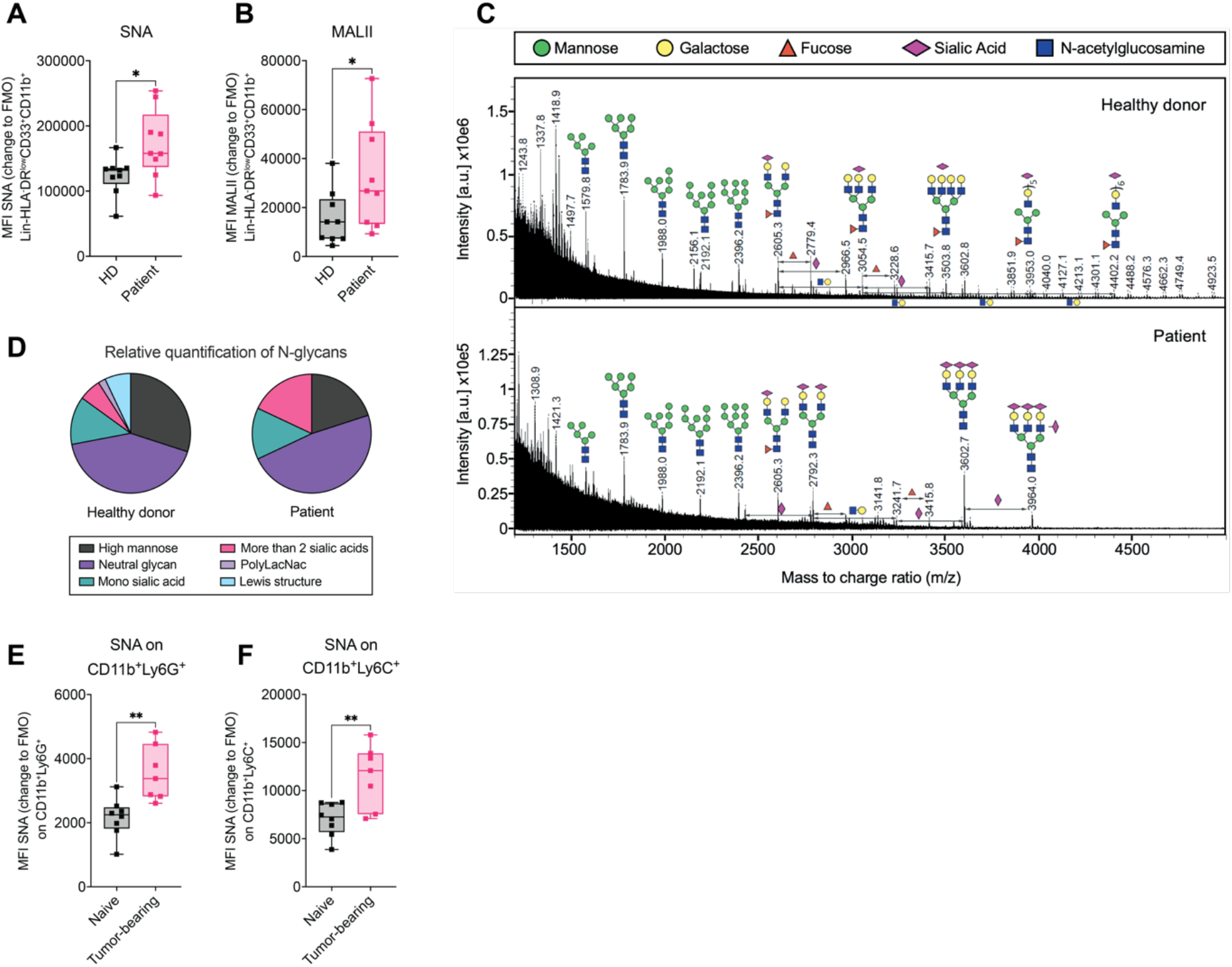
Myeloid cells in cancer are highly sialylated. **(A)** MFI of SNA or **(B)** MALII gated on PB-derived Lin^-^HLADR^low^CD33^+^CD11b^+^ cells from lung cancer patient and healthy controls. MFI is shown as change to FMO and was determined by flow cytometry. *N=8-13 donors with at least N=2 experiments.* **(C)** MALDI-TOF mass spectra (m/z 1200–5000) of N-glycans isolated from CD33^+^ cells of healthy donor and lung cancer patient derived from fresh blood. The N-glycans were released by PNGaseF and permethylated prior to MALDI-TOF-TOF profiling. Main structures are depicted above the corresponding peaks. Assignments are based on composition and knowledge of biosynthetic pathways. All molecular ions are [M + Na]^+^. Residues above a bracket have not had their location unequivocally defined. **(D)** Relative quantification of N-Glycans detected in cancer patient and healthy donor derived CD33^+^ cells from **(C)**. *N=1*. **(E)** Fresh blood from B16F10 tumor bearing mice and naïve wildtype mice was collected at day 14 after tumor inoculation and analyzed for SNA gated on **(E)** CD45^+^CD11b^+^Ly6C^+^ or **(F)** CD45^+^CD11b^+^Ly6G^+^ cells. MFI is shown as change to FMO. *7-8 mice per group with N=2 experiments*. Data are presented as mean. Error bar values represent SD. Two-tailed unpaired Student’s t test was used. *P<0.05, **P<0.01, ***P<0.001, and ****P<0.0001.

We further studied the expression of ligands on the surface of murine myeloid cells by lectin staining. Similar to the human setting, increased levels of SNA were detected on both, CD11b^+^Ly6G^+^ and CD11b^+^Ly6C^+^ populations in the blood of tumor-bearing mice compared to naïve littermates (2 E, F). No significant differences could be observed on MALII or PNA level (Supp.2 B, C). This data shows differential expression of sialoglycans on cancer-associated suppressive myeloid cells in humans and mice.

### 3. Depletion of Siglec-E on myeloid cells prolongs survival

To further investigate the role of Siglec-E on myeloid cells in the TME, Siglec-E^loxP^ mice were crossed with LysMCre to specifically target Siglec-E on LysM expressing cells in mice (SigE^ΔLysM^). SigE^ΔLysM^ were compared to Siglec-E wildtype (SigE^WT^) littermates in terms of tumor growth, survival and immune infiltration upon subcutaneous tumor injection (3 A). As we were focusing on suppressive myeloid cells, models of cancer-induced emergency myelopoiesis were used, including B16F10 melanoma and EL4 lymphoma syngeneic tumor models (22). During emergency myelopoiesis, myeloid cells are rapidly increased which leads to accumulation of immature, suppressive cells including MDSCs.

**Figure 3:**
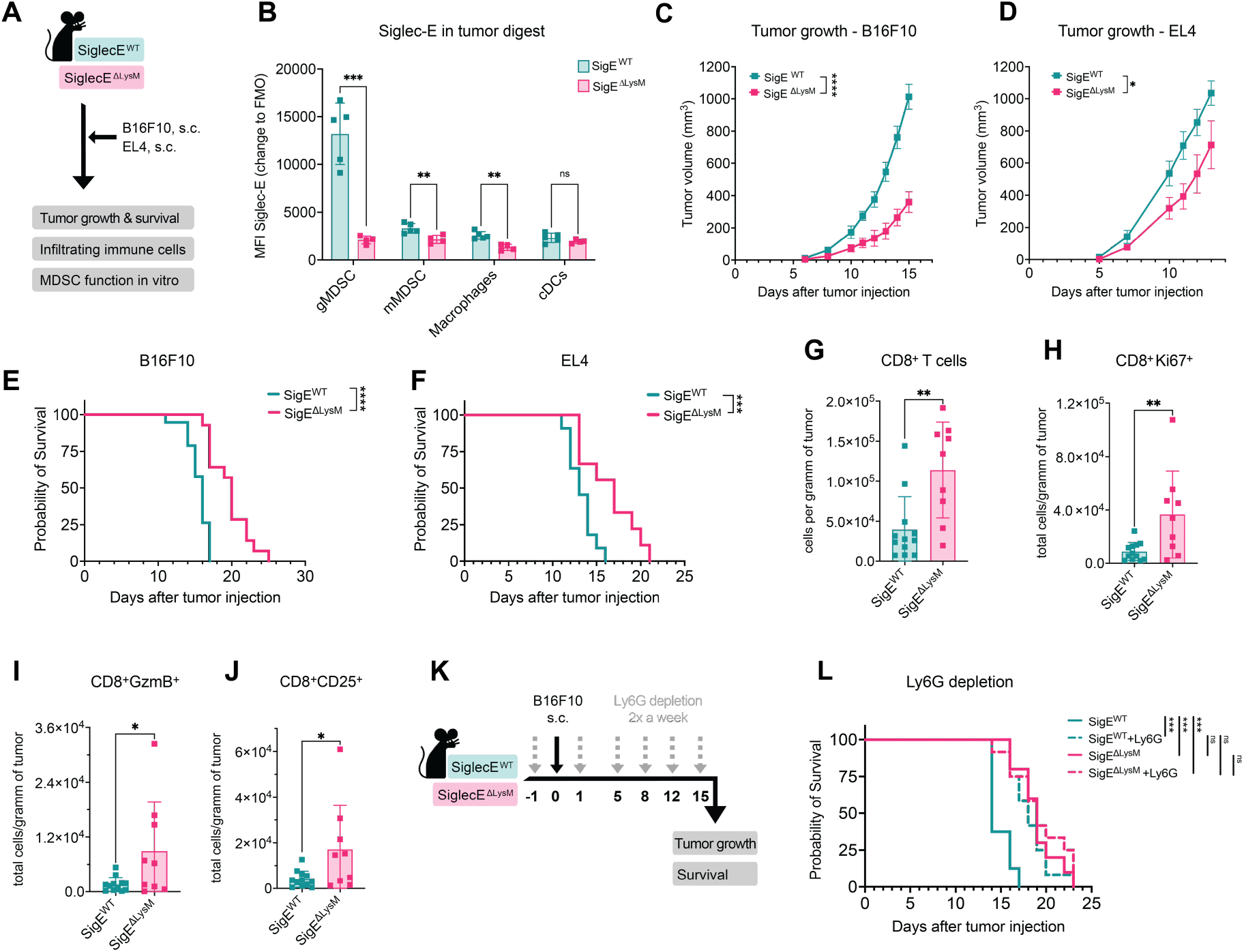
Siglec-E depletion on myeloid cells decreases tumor growth in mice. **(A)** Experimental setup: Siglec-ExLysM-Cre mice (SigE^ΔLysM^) and Siglec-E WT (SigE^WT^) littermates were subcutaneously injected with B16F10 or EL4 cells. Tumor growth, probability of survival, tumor immune cell infiltration and suppressive capacity of Gr1^+^CD11b^+^ *in vitro* were analyzed. **(B)** MFI of Siglec-E expression was assessed on myeloid cells in tumor digest at endpoint of SigE^ΔLysM^ mice and SigE^WT^ littermates. MFI of Siglec-E is shown as a change to FMO. Cells were identified as gMDSCs (CD45^+^CD11b^+^Ly6G^+^), mMDSCs (CD45^+^CD11b^+^Ly6C^+^), macrophages (CD45^+^CD11b^+^F4/80^+^), dendritic cells (DCs) (CD45^+^CD11c^+^MHCII^+^F4/80^-^). *N=4-5 mice per group*. **(C)** Tumor growth from pooled data of B10F10 and **(D)** EL4 subcutaneously injected mice. *N=9-12 mice per group from at least 2 independent experiments*. **(E)** Kaplan-Meier survival curves from pooled data of mice injected subcutaneously with B16F10. *N=9-12 mice per group from at least 2 independent experiments*. **(F)** Kaplan-Meier survival curves from pooled data from 2 experiments injected with EL4. *N=9 mice per group* **(G)** B16F10 tumors at endpoint **(C)** were digested and analyzed by flow cytometry. Intratumoral CD8^+^ cells (CD45^+^CD19^-^NKp46^-^CD3^+^CD8^+^), **(H)** Ki67^+^CD8 T cells, **(I)** GranzymeB^+^CD8 T cells^+^ (GzmB^+^), **(J)** CD25^+^CD8 T cells were quantified as cells per gram of tumor at the endpoint of the experiment. Pooled data from 2 independent experiments. *N=9-12 mice per group.* **(K)** Experimental setup: Depletion of Ly6G positive cells using depletion antibody in SigE^ΔLysM^ mice and SigE^WT^ littermates bearing B16F10 tumors. Mice were injected up to 6 times (grey arrow) with Ly6G depletion antibody starting 1 day before subcutaneous B16F10 tumor injection (black arrow). Tumor growth and survival were monitored. **(L)** Kaplan-Meier survival curves from pooled data from 2 independent experiments. *N=9-11 mice per group*. Data are presented as mean. Error bar values represent SD. Two-tailed unpaired Student’s t test or multiple unpaired t-tests (B) was used. For survival analysis, log-rank test was used followed by Šidák correction for multiple comparisons. Tumor growth was compared by mixed-effects analysis followed by Bonferroni’s multiple comparisons test. *P<0.05, **P<0.01, ***P<0.001, and ****P<0.0001.

Subcutaneous injection of both tumor cell lines resulted in increased numbers of CD11b^+^Ly6G^+^ and CD11b^+^Ly6C^+^ in the spleen of tumor-bearing mice compared to naïve littermates (Supp.3 A, B). Additionally, high numbers of MDSCs were found within B16F10 and EL4 tumors, making them suitable models to study the effect of Siglec-E in myeloid-driven tumors (Supp.3 C). To confirm deletion of Siglec-E in our model, Siglec-E expression was accessed by flow cytometry on myeloid cells from tumor digests and spleens of EL4 and B16F10 tumor-bearing mice (3 B, Supp.3 D, E). Siglec-E was significantly decreased on gMDSCs, mMDSCs and macrophages in the tumor and spleen of SigE^ΔLysM^ compared to SigE^WT^ mice. Siglec-E was the highest expressed on gMDSCs followed by mMDSCs and macrophages in wildtype mice within the tumor and the spleen (3 B, Supp.3 D,E) as observed before (1 E, F). Deletion of Siglec-E on myeloid cells resulted in prolonged survival and decreased tumor growth of SigE^ΔLysM^ mice compared to SigE^WT^ littermate mice in B16F10 and EL4 tumor models (3 C-F). We observed increased CD8^+^ T cell infiltration in SigE^ΔLysM^ mice associated with an increased number of proliferation and expression of functional T cell markers including Granzyme B (GzmB), Ki67 and CD25 (3 G-J). To avoid cancer model-dependent effects, we confirmed our findings using EL4 lymphoma tumor cells. Increased CD8^+^ T cell infiltration was also found in EL4 bearing mice strengthening the importance of our findings (Supp.3 F-I).

As the highest expression of Siglec-E was found on gMDSCs, we hypothesized that Siglec-E depletion mainly affects gMDSCs in our model. To test this hypothesis, Ly6G depletion was performed to eliminate Ly6G-expressing gMDSCs in SigE^ΔLysM^ mice and SigE^WT^ littermates upon B16F10 tumor cell injection (3 K). Depletion of gMDSC in SigE^WT^ mice prolonged their survival and tumor growth but did not affect SigE^ΔLysM^ mice lacking Siglec-E on myeloid cells (3 L, Supp.3 J). Thus, Siglec-E expression on gMDSCs was likely involved in tumor progression *in vivo*. The numbers of myeloid cells within the tumor and spleen were not altered (Supp. 3 K-N) suggesting a qualitative change rather than a quantitative change of myeloid cells upon Siglec-E depletion, which possibly resulted in a less suppressive TME leading to effector T cell infiltration. These results show that Siglec-E on myeloid cells inhibits tumor growth in different murine tumor models, mainly due to Siglec-E expression on gMDSCs.

### 4. Siglec-E and sialoglycans shape the immunosuppressive capacity of murine MDSCs

Although MDSCs are involved in various pro-tumorigenic mechanisms, the key feature remains their ability to inhibit T cell response (17). To test whether the suppressive function of MDSCs lacking Siglec-E is impaired, MDSCs were isolated by CD11b^+^Gr1^+^ negative selection from the spleen of B16F10-tumor bearing mice and their suppressive capacity was tested against highly stimulated naïve T cells *in vitro* (4 A). Stimulation of T cells by IL-2 and anti-CD3/28 led to high proliferation of T cells (4 B). MDSCs from SigE^WT^ mice highly suppressed CD8 T cell proliferation but SigE^ΔLysM^ MDSCs were significantly less suppressive shown by an increased T cell proliferation (4 C, D). This suggests that the reduced suppressive function of MDSCs lacking Siglec-E could generate a less suppressive TME *in vivo* leading to increased T cell infiltration and prolonged survival that is observed in SigE^ΔLysM^ mice.

**Figure 4:**
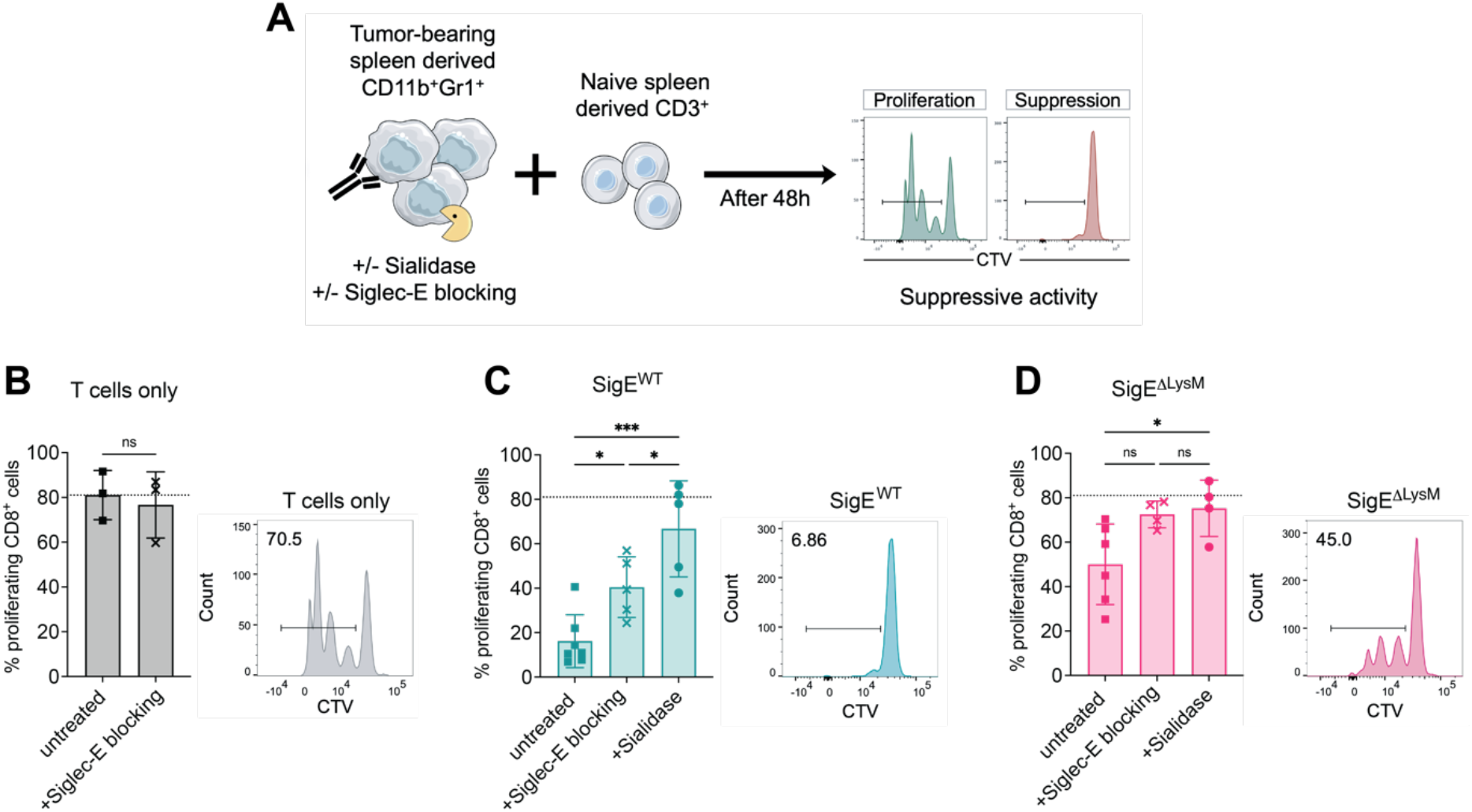
Reduced suppressive function of MDSCs lacking Siglec-E and upon sialidase or Siglec-E blocking antibody treatment. **(A)** Experimental setup to assess the suppressive capacity of Gr1^+^CD11b^+^ (MDSCs) cells against naïve CD3^+^ (T cells) cells. MDSCs were isolated from spleens of B16F10 tumor-bearing SigE^ΔLysM^ mice and SigE^WT^ littermates. T cells were isolated from naïve littermates, stained with CTV and co-cultured with MDSCs for 48 hours in the presence of aCD3, aCD28 and IL-2. MDSCs were used immediately or pretreated with sialidase. Siglec-E blocking antibody was added to the co-cultures as indicated. **(B)** Percentage of proliferation of CD8^+^ T cells co-cultured without MDSCs, **(C)** with MDSCs from SigE^WT^ mice or **(D)** MDSCs from SigE^ΔLysM^ mice. Exemplary results for each untreated condition are shown on the right. Pooled data from at least 2 independent experiments*. N=3-7 mice per group from N=3 experiments.* Data are presented as mean. Error bar values represent SD. Two-tailed unpaired Student’s t test or multiple unpaired t-tests (C, D) was used. *P<0.05, **P<0.01, ***P<0.001, and ****P<0.0001.

To further evaluate if suppressive MDSCs can be altered by interfering with the Siglec-sialoglycan axis, we either added Siglec-E blocking antibody to the co-cultures or pretreated MDSCs with bacterial sialidase to reduce the level of both α2,3- and α2,6-sialoglycans on the surface of MDSCs. Using a Siglec-E blocking antibody, we could decrease the suppression by SigE^WT^ MDSCs but no significant difference was observed by blocking Siglec-E on SigE^ΔLysM^ MDSCs or T cells alone (4 B-D). Pretreatment of MDSCs with sialidase strongly reduced MDSC suppressive activity of both, SigE^ΔLysM^ and SigE^WT^-derived MDSCs (4 C, D). This data shows that sialoglycan ligands and Siglec-E on murine MDSCs are important players in the suppressive effect of MDSCs against murine T cells.

To further address the effect of sialidase treatment and lack of Siglec-E *in vivo*, we generated B16F10 cells stably expressing *H1N1* viral sialidase (B16F10-sia) and compared tumor growth with B16F10 wildtype in SigE^ΔLysM^ and SigE^WT^ mice (Supp.4 A+B). In accordance with *in vitro* experiments, sialidase expression highly decreased tumor growth in SigE^ΔLysM^ and SigE^WT^ mice (Supp.4 C+D). Mice injected with B16F10-sia that additionally were SigE^ΔLysM^ showed the highest survival benefit and resulted in 50% tumor free-survival. However, SigE^WT^ mice injected with B16F10-sia eventually reached tumor endpoint. Taken together, Siglec-E as well as sialoglycan ligands on murine MDSCs are involved in the suppression of CD8^+^ T cells.

### 5. Sialoglycans modulate the generation of human suppressive myeloid cells generated *in vitro*

Next, we were wondering if targeting the Siglec-sialoglycan axis on human MDSCs affects their suppressive capacity. To test this, we used an *in vitro* model to generate suppressive tumor-educated myeloid-derived CD33^+^ cells further referred to as MDSC-like cells by adapting the protocol from Lechner et al. (23) (5 A). Using lung adenocarcinoma A549 and cervix adenocarcinoma HeLa cell lines, we generated highly immunosuppressive MDSC-like cells from fresh peripheral blood mononuclear cells (PBMCs) that were able to decrease CD8^+^ T cell proliferation in an effector:target (E:T) ratio-dependent manner (Supp.5 A). This model was used to test the role of the Siglec-sialoglycan axis during MDSC generation and function with suppression of autologous T cells as a functional read-out similar to the previous murine studies.

**Figure 5:**
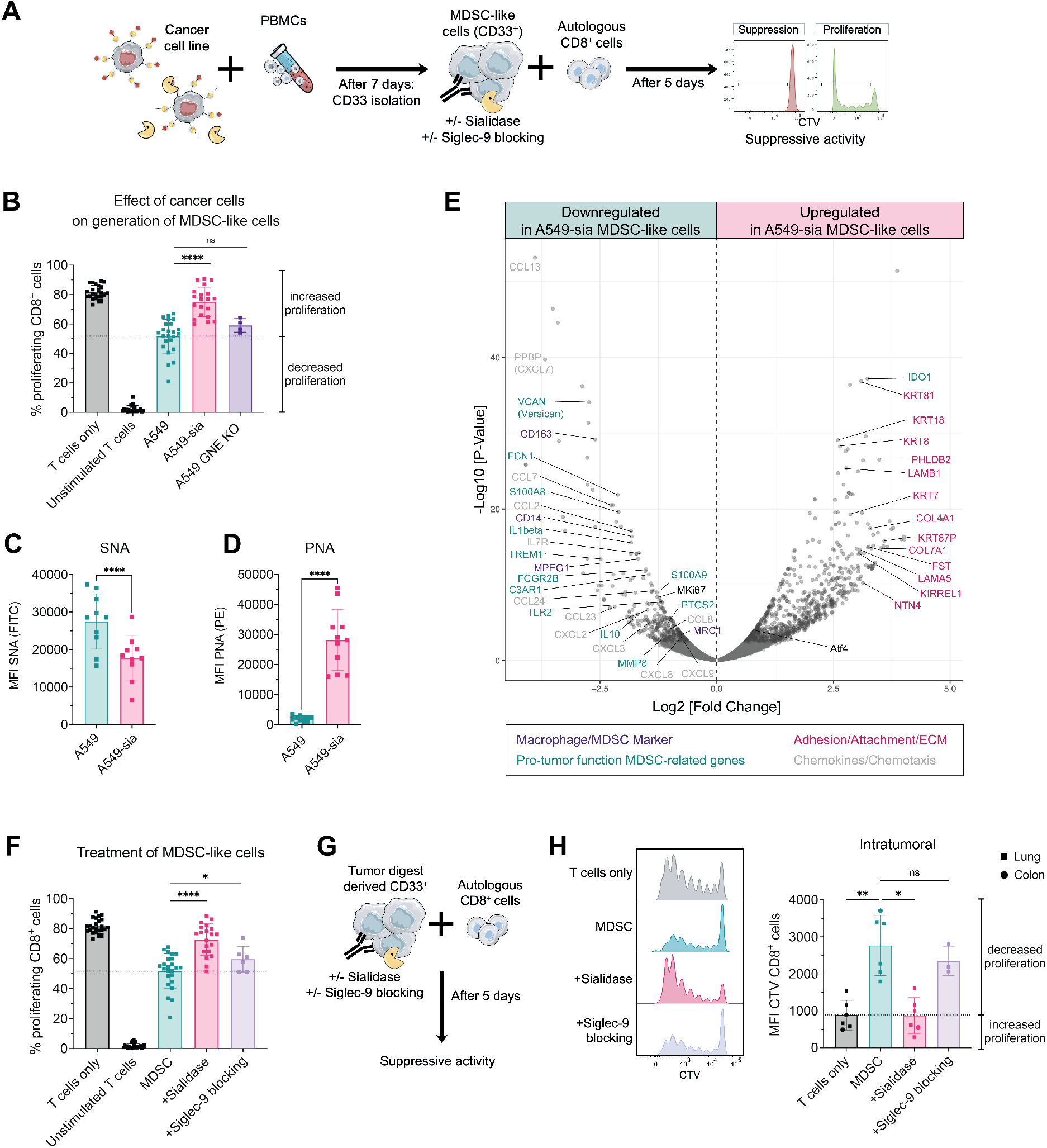
Targeting sialoglycans and Siglec-9 on suppressive human CD33^+^ cells attenuates their function. **(A)** Experimental setup to generate suppressive myeloid cells *in vitro*. Fresh PBMCs were isolated from buffy coats from healthy donors and co-cultured with indicated cancer cells lines at a ratio of 100:1. On day 7, CD33^+^ cells were isolated by magnetic positive selection and their suppressive capacity was assessed against autologous CD8^+^ T cells. Suppressive CD33^+^ cells were immediately used, pretreated with sialidase or Siglec-9 blocking antibody was added to the co-culture. CD8^+^ T cells were stained with CTV and stimulated by addition of IL-2 and anti-CD3/28 microbeads. After 5 days, CD8^+^ T cell proliferation was assessed by FACS **(B)** Percentage of proliferating CD8^+^ cells upon co-culture with indicated suppressive CD33^+^ cells. Suppressive myeloid cells were generated using A459, A549 stably expressing sialidase (A549-sia) or A549 GNE KO cancer cell lines. N=4-24 donors in at least n=3 experiments **(C)** Lectin staining was performed on suppressive CD33^+^ cells on day 7 of the experiment assessing SNA and **(D)** PNA. *N=10 donors from N=5 experiments.* **(E)** Volcano plot of differentially expressed genes of suppressive CD33^+^ cells generated with A549 or A549-sia cancer cell lines. CD33^+^ cells were isolated on day 7 and processed for bulk RNA Sequencing. *N=4 per group*. **(F)** Percentage of proliferating CD8^+^ cells upon co-culture with suppressive CD33^+^ cells generated by A549 co-culture. CD33^+^ cells were used immediately, pretreated with sialidase or Siglec-9 blocking antibody was added to the co-culture. *N=6-24 donors from at least n=4 experiments* **(G)** Assay setup to test the suppressive capacity of tumor-digest derived CD33^+^ cells. CD33^+^ cells were isolated freshly from tumor digest of lung or colon cancer patients, pretreated with sialidase or Siglec-9 blocking antibody was added to the co-cultures. CD8^+^ cells were isolated from fresh PBMCs and stained with CTV. The suppressive activity was assessed on day 5 by flow cytometry. **(H)** MFI of CTV staining of CD8^+^ cells is shown upon co-culture with tumor-derived CD33^+^ cells. MDSCs were untreated, pretreated with sialidase or Siglec-9 blocking antibody was added to the co-culture. Exemplary results for each condition are shown on the left. *N=3-6*. Data are presented as mean and error bar values represent SD. Paired t-test or one-way ANOVA (H) was used. *P<0.05, **P<0.01, ***P<0.001, and ****P<0.0001.

To determine the role of the glycosylation of cancer cells during generation of MDSC-like cells, we compared the potential of parental A549, A549 cells expressing sialidase (A549-sia) and A549-GNE Knockout (KO) cell lines to generate MDSC-like cells. A549-GNE K cells have a deficiency of UDP-N-acetylglucosamine 2-epimerase/N-acetylmannosamine kinase (GNE), a key enzyme of sialic acid biosynthesis leading to decreased sialoglycan expression (24). A549-sia stably express membrane-bound viral sialidase, which cleaves α2,3- and α2,6-sialic acid from the surface of A549 cells and also surrounding cells. To test the sialidase activity, A549 cells were stained for lectins indicating effective desialylation by an increase in PNA and decrease in SNA and MALII levels (Supp.5 B). Generation of MDSC-like cells with parental cancer cells as well as GNE KO cells resulted in strong suppressive capacity. In contrast, co-culture with sialidase-expressing cancer cells induced a significantly less suppressive phenotype shown by an increased T cell proliferation (5 B). To avoid a cell line-specific effect, we used HeLa and HeLa cells expressing sialidase (HeLa-sia) and observed similar results (Supp.5 D). To test the effect of sialidase expression on MDSC-like cells, we assessed the lectin levels of MDSC-like cells in A549-sia co-cultures. Co-culture with A549-sia led to desialylation of MDSC-like cells as shown by a decrease in SNA and MALII levels as well as an increase in PNA compared to MDSC-like cells generated with parental cancer cell lines (5 C+D, Supp.5 C). These findings suggest that the sialoglycan levels on MDSC-like cells are important for their suppressive function against T cells, but the level of sialoglycan ligands on cancer cell lines does not impact their suppressive potential. This indicates an interaction of Siglecs and sialoglycan ligands on the surface of MDSC-like cells rather than *cis* interaction with cancer cells.

To better understand the differences of *in vitro* generated MDSC-like cells and the role of sialoglycans during MDSC generation, transcriptomics analysis was performed by bulk RNA sequencing of MDSC-like cells generated with A549 and A549-sia cancer cell lines. Suppressive myeloid cells created with A549-sia resulted in a downregulation of various functional markers previously described as MDSC markers or pro-tumor function MDSC-related genes on RNA level including S100A8/9, PTGS2, IL10 and IL1beta (17,25)(5 E, Supp.5 E). Additionally, generation of MDSC-like cells by A549-sia co-culture significantly inhibited chemokine and chemotaxis molecules on RNA level including CCL2, CCL13, CXCL7 and CXCL2 and led to an increase in many genes involved in adhesion and attachment. These results show that desialylation of MDSC-like cells during their generation leads to downregulation of MDSC-functional markers on RNA level and decreased suppressive capacity *in vitro*.

### 6. Sialidase treatment and Siglec-9 blocking attenuate the suppressive activity of myeloid cells

Next, we wanted to address the effect of sialidase treatment and Siglec-9 blocking as a therapeutic approach to treat *in vitro* generated human suppressive myeloid cells. To this end, we generated MDSC-like cells as described in the previous section and pretreated them either with bacterial and viral sialidase to cleave surface sialoglycan ligands or added Siglec-9 blocking antibody to the co-cultures (5 A+F, Supp. 5 F). Sialidase pretreatment and blocking of Siglec-9 with an antibody resulted in a significant decrease of the suppressive capacity of MDSC-like cells against autologous T cells (5 F, Supp. 5 F). Successful desialylation of cells by sialidase treatment was demonstrated by a significant increase in PNA staining (Supp. 5 G). Similar results were obtained using HeLa-generated MDSC-like cells (Supp. 5 H).

To further corroborate our findings, we used cancer patient-derived CD33^+^ cells from primary tumor cell suspensions together with autologous CD8^+^ T cells isolated from PBMCs (5 G). The addition of tumor-derived CD33^+^ cells from colon and lung cancer patients could significantly decrease the proliferation of CD8^+^ T cells (5 H). Pretreatment of suppressive myeloid cells with sialidase led to significant reduction of their inhibitory effect on CD8^+^ T cell proliferation (5 H). Although a trend towards downregulation of suppressive capacity was observed, addition of Siglec-9 blocking antibody did not lead to significant changes compared to untreated CD33^+^ cells, suggesting that other sialic acid-binding receptors including other Siglec receptors could be involved. Our experiments using human MDSC-like cells and intratumoral patient-derived suppressive CD33^+^ cells support our finding that the interactions of Siglec receptors with cell surface sialoglycan ligands on MDSCs can regulate their suppressive potential which is an interesting target to attenuate MDSC’s suppressive function.

### 7. Reduction of sialoglycan ligands on MDSCs reduces immune-inhibitory CCL2 production and enhances anti-cancer immunity

Suppressive myeloid cells are involved in various pro-tumorigenic mechanisms which can be mediated by the production of suppressive cytokines or chemokines (26). To better understand the underlying mechanism responsible for changes of MDSC function influenced by Siglec-sialoglycan interactions, we analyzed the cytokines and chemokines in MDSC-T cell co-culture supernatants by ELISA. By checking murine co-culture supernatants (from 4 B-D), we found various cytokines in co-cultures compared to T cells alone, including CCL2, IL1β, IL-6 and IL-10 (Supp. 6 A). CCL2 was highly increased in supernatants of suppressive SigE^WT^ supernatants compared to SigE^ΔLysM^ and sialidase treated conditions (6 A, Supp. 6 A). Furthermore, the suppressive capacity of MDSCs strongly correlated with CCL2 detected in the supernatant, indicating a relevant role of CCL2 in MDSC function (Supp. 6 B).

**Figure 6:**
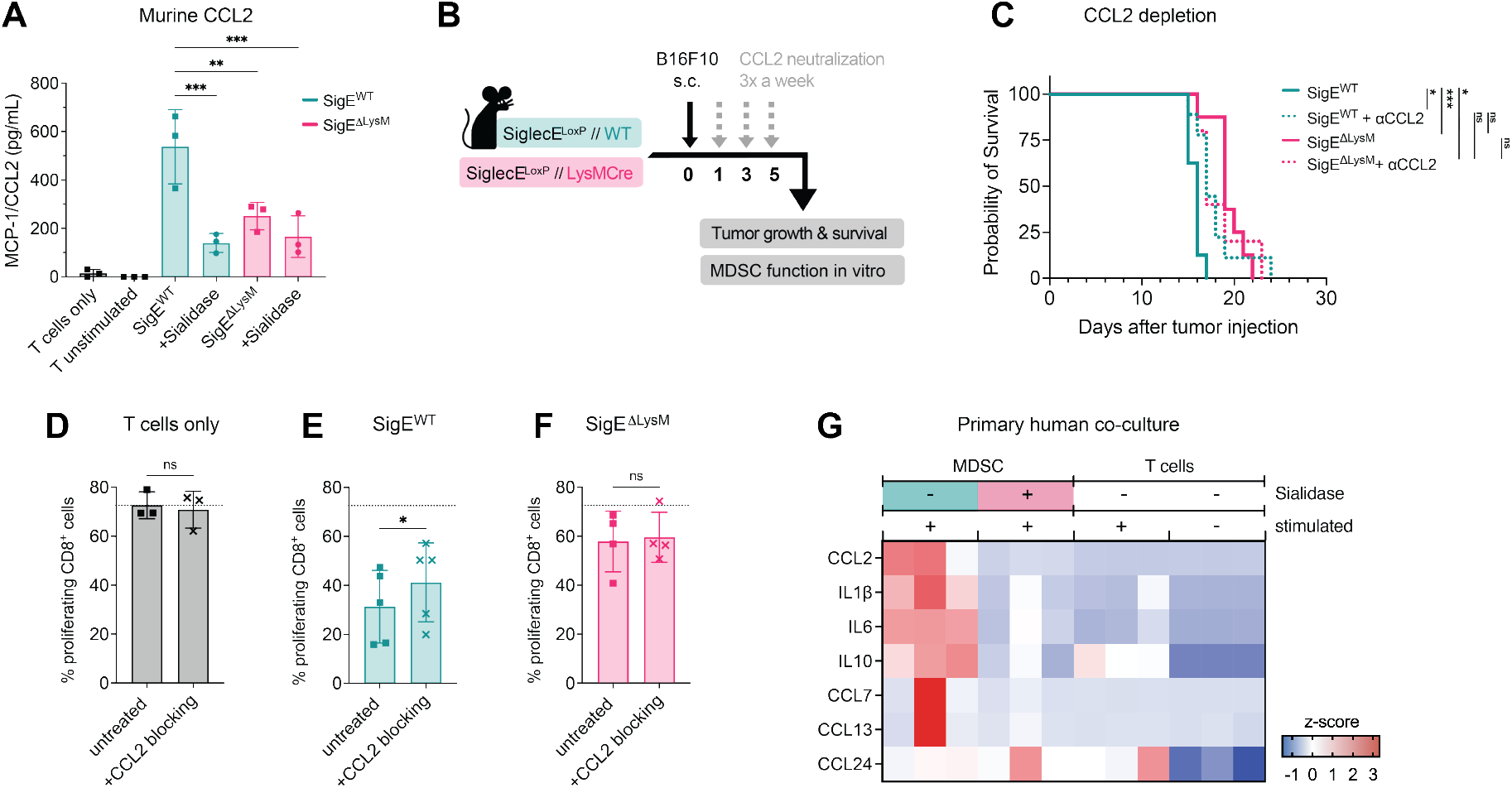
CCL2 is involved in T cell suppression via Siglec-sialoglycan axis on suppressive myeloid cells. **(A)** MCP-1/CCL2 found in the supernatant of murine MDSC:T cell co-cultures at endpoint of the experiment from figure 4 B-D. MDSCs were untreated, pretreated with sialidase or Siglec-E blocking antibody was added to the co-culture. *N=3 donors per group*. **(B)** Experimental setup: Neutralization of CCL2 using neutralization antibody in SigE^ΔLysM^ mice and SigE^WT^ littermates bearing B16F10 tumors. Mice were injected with CCL2 neutralization antibody up to 3 times a week (grey arrow) starting 1 day after subcutaneous B16F10 tumor injection (black arrow). Tumor growth and survival were monitored and suppressive capacity of MDSCs was analyzed *in vitro*. **(C)** Kaplan-Meier survival curves from pooled data from 2 independent experiments. *N=5-8 mice per group.* **(D)** Suppressive capacity of MDSCs against naïve T cells. Percentage of proliferation of CD8^+^ T cells co-cultured without MDSCs, **(E)** with MDSCs from SigE^WT^ mice or **(F)** MDSCs from SigE^ΔLysM^ mice with or without CCL2 blocking antibody. *N=3-5 mice from N=3 experiments* **(G)** Cytokine expression found in the supernatant of human primary CD33^+^:CD8^+^ cell co-cultures at endpoint of the experiment from figure 5 H. CD33^+^ cells were untreated or pretreated with sialidase. Z-scores were calculated for each cytokine and are shown on a color scale from blue to red. *N=3 donors per group*. Data are presented as mean and error bar values represent SD. Paired t-test was used. *P<0.05, **P<0.01, ***P<0.001, and ****P<0.0001.

CCL2 is widely described in the context of MDSCs and can act as a chemoattractant which is involved in the migration of myeloid cells and contributes to intratumoral MDSC accumulation (27). Apart from its role as a chemoattractant, CCL2 facilitates immunosuppression of T cells by regulating suppressive functions of MDSCs via STAT3 in colorectal cancer (28) and is not only expressed by cancer cells, but also by TAMs and MDSCs (29,30). To further evaluate the role of CCL2 as a mediator of MDSC suppression, the effect of CCL2 neutralization on tumor growth in SigE^ΔLysM^ and SigE^WT^ mice was addressed (6 B). Importantly, CCL2 neutralization *in vivo* lead to a prolonged survival in SigE^WT^ mice, but did not significantly alter the survival of SigE^ΔLysM^ indicating an involvement in Siglec-E signaling on myeloid cells (6 C). To test whether CCL2 was directly involved in the suppressive capacity of MDSCs, we analyzed the effect of CCL2 blocking antibody on MDSCs function against T cells *in vitro* (6 D-F). In accordance with the *in vivo* results, CCL2 blocking significantly decreased the suppressive function of SigE^WT^ MDSCs (6 E), but did not impact SigE^ΔLysM^ MDSCs (6 F).

Similar to murine cell co-culture, the CCL2 transcript was significantly downregulated in conditions when MDSC-like cells were generated with A549-sia (5 E). To further investigate the effect of sialidase treatment on chemokine and cytokine expression by human MDSCs, cytokine levels were measured in primary human co-culture from intratumoral suppressive myeloid cells (from 5 H). As observed in mice, high amounts of CCL2 were detected in supernatants of suppressive myeloid cells but pretreatment of primary human intratumoral CD33^+^ cells with sialidase showed significantly diminished CCL2 secretion (6 G, Supp. 6C). Additionally, high levels of IL1β, IL-6 and IL-10 were detected in suppressive CD33^+^ cells supernatants (6 G). Sialidase treated CD33^+^ cells and T cells alone showed low to no expression. These results suggest that interactions of cell surface sialoglycan ligands with Siglec receptors induce a suppressive phenotype in myeloid cells that inhibit sufficient anti-cancer immunity by secretion of immune-inhibitory CCL2.

## Discussion

Although the Siglec-sialoglycan axis is gaining attention as a potential glyco-immune checkpoint in cancer, little is known about the expression and function of Siglecs and sialoglycan ligands on suppressive myeloid cells (10). Here, we show that targeting cell surface sialoglycan ligands and Siglec receptors on MDSCs can decrease their suppressive capacity by downregulation of cytokines and chemokines mainly via CCL2.

Previous preclinical studies have demonstrated anti-cancer effects of blocking Siglec receptors and/or sialidase treatment on various immune cell types including T cells, NK cells and myeloid cells such as TAMs, resulting in a TME permissive towards successful cancer immunotherapy (6,11,12,31–34). Here, we advance the understanding how Siglec-sialoglycan interactions on myeloid cells can shape an immunosuppressive environment via secretion of inhibitory CCL2 in the context of cancer across different human and murine models. Although we observed a strong effect of Siglec-E deletion and blocking on decreasing the suppressive capacity of suppressive myeloid cells in mice, blocking of a single inhibitory Siglec receptor on suppressive myeloid cells by Siglec-9 blocking antibody resulted in a less pronounced effect in human cell culture models. This could potentially be explained by the fact that human suppressive myeloid cells express different, maybe redundant inhibitory CD33-related Siglec receptors compared to murine suppressive myeloid cells that mainly express Siglec-E in cancer. It is possible that other Siglec family members like Siglec-5, Siglec-7 or Siglec-10 are involved in modulating the suppressive capacity of human MDSCs and that their importance might be variable or interchangeable between different tumors (12). Sialidase treatment might be able to circumvent this by cleaving ligands for multiple Siglec receptors. A strong effect of sialidase pretreatment on the suppressive capacity of myeloid cells was observed across all assays which supports this hypothesis. A first-in-human trial using a human bi-sialidase as cancer immunotherapy against solid tumors showed tolerability and desialylation of immune cells in the peripheral blood (35) (NCT05259696). Additional work will be needed to investigate the effect of sialidase treatment on MDSCs in this setting, but it seems encouraging to affect various players generating a suppressive TME including MDSCs and TAMs (11).

Reduction of sialoglycans on cancer cells did not affect the suppressive capacity of MDSC-like cells, but strong effects were observed upon constant expression of sialidase by cancer cells resulting in a desialylation of MDSC-like cells. Similarly, the suppressive capacity of MDSCs was strongly reduced in human and murine MDSC:T cell co-culture models upon pre-treatment of MDSCs with sialidase. This data supports the hypothesis that interactions of Siglec receptors with sialoglycan ligands on myeloid cells plays a more important role as *cis* interaction than the previously propagated *trans* interactions of Siglec receptors with sialoglycan ligands on cancer cells. Expression of *cis*-ligands on various immune cells was described before and proposed as possible mechanism on MDSCs and immature DCs (16,36). However, further studies are needed to investigate whether Siglecs and sialoglycan ligands on MDSCs interact *cis* on the same cell or *trans* between neighboring MDSCs.

Most Siglec receptors are classified as inhibitory receptors, which harbor tyrosine-based signaling motifs like ITIM domains that can recruit and activate tyrosine phosphates including SHP-1 and SHP-2 (5). We identified the interaction of sialoglycan ligands and Siglecs on MDSCs as stimulus for MDSCs leading to the release of suppressive cytokines (CCL2, IL-6, IL-10). Blockade of this interaction resulted in a decreased suppressive function. In line with our findings, others demonstrated activating signaling of macrophages and monocytes upon Siglec-9 engagement via the MEK/ERK pathway (33). Additionally, binding of CD33/Siglec-3 by the S100A9 family on MDSCs is involved in MDSC expansion and accumulation resulting in a release of suppressive cytokines (37). Therefore, it seems like the expression of Siglecs and sialoglycan ligands is highly context and cell type specific and can on one hand cause “classical” engagement and ITIM domain signaling and on the other hand result in a positive feedback loop maintaining signaling of suppressive myeloid cells. However, it is not yet clear what exact type of sialoglycan ligands is involved and further studies are needed to understand the exact mechanism how sialoglycans can support immunosuppressive properties of suppressive myeloid cells in cancer. Moreover, it also remains unclear what role other lectins play after using sialidase and exposing lactosamine residues that could again bind to other immunomodulatory lectins including galectins (38).

Additionally, we identified CCL2 as an important immune-inhibitory chemokine released upon interactions of inhibitory Siglec receptors and sialoglycan ligands on suppressive myeloid cells. Previously, CCL2 was described to impact the secretion of effector molecules and to contribute to T cell suppression via STAT3 signaling in MDSCs (27,28). Additionally, myeloid cells express high levels of CCR2 which is a promising target to interfere with MDSC migration to the TME and can also express CCL2 themselves (13,29,30). Therefore, it is not surprising that we found a strong association between the suppressive capacity of MDSCs and CCL2 and that blocking of CCL2 led to an improved T cell proliferation. Nevertheless, we are the first once to propose a linkage of the Siglec-sialoglycan axis on MDSCs with CCL2 expression. Further investigation is needed to better understand the molecular mechanism and role of other cytokines involved including IL1β, IL-6 and IL-10.

Our study contains some limitation. By using human CD33^+^ cells and murine LysM-Cre models, we target a variety of myeloid cells and it would be desirable to specifically target each of the suppressive myeloid cell subtypes to unveil their individual contribution to immunosuppression. However, suppressive myeloid cells are closely related and recent publications utilizing in-depth transcriptional, biochemical and phenotypical characterization reveal the high complexity and plasticity of those cells (13,39–41). A clear distinction and definition by phenotype as well as the functional relevance of subtypes of myeloid cells is still lacking. Additionally, our assays focus on the suppressive function of MDSCs against T cells, which is described as gold standard (17). Interaction of Siglecs and sialoglycans on MDSC may also have additional functions on other immune cells which need to be addressed.

Taken together, cancer-associated suppressive myeloid cells express high levels of inhibitory Siglec receptors and cognate sialoglycan ligands inducing an immunosuppressive phenotype. We also identify CCL2 as a major inhibitory mediator of this effect. Blocking of the Siglec-sialoglycan axis using sialidase or another broader approach targeting different Siglec receptors could potentially render an immunosuppressive TME permissive for cancer immunotherapy including immune checkpoint inhibition. Targeting sialoglycans or Siglec receptors could therefore be used to treat cancers with a significant infiltration of suppressive myeloid cells.

## Acknowledgments

We thank Mélanie Buchi for her technical support and all members of the Cancer Immunology and Cancer Immunotherapy Laboratory of the Department of Biomedicine for helpful discussions. We are grateful to all patients providing material. Furthermore, we want to thank the Flow Cytometry, Genomics and Animal core facilities of the Department of Biomedicine for their support. This work was supported by funding from the Goldschmidt-Jacobson Foundation, the Swiss National Science Foundation (SNSF Grant No. 310030-184720), the Schoenmakers-Müller Foundation and the Cancer League of Basel (KlbB).

## Author contributions

R.W., H.L. and N.R.M. planned the project and experimental design. R.W., A.Zingg, E.C., C.W.L., P.H. and L.D. performed and analyzed experiments. R.W., H.L. and N.R.M interpreted the results. A.B. processed and analyzed RNA data and performed data depositions. R.W., H.L. and N.R.M. wrote the manuscript. R.W., E.C., A.B., C.W.L., A.Zingg, P.H., A.Zippelius, H.L. and N.R.M. reviewed the manuscript. D.L. provided clinical samples. H.L. and A.Zippelius collected the clinical data and ethical board approvals. R.W. designed the graphical summary. All authors read and approved the final manuscript.

## Materials and Methods

### Cell lines

HeLa and B16F10 cell lines were obtained from ATCC. A549, HeLa and EL4 were kindly provided by Zippelius Lab and HEK293T by the Bentires Lab, both from the Department of Biomedicine, Basel. *H1N1* viral sialidase expressing cell lines, A549-sia, HeLa-sia, B16F10-sia as well as EL4 GFP cells were generated by lentiviral transduction as described below. A549-GNE KO cells were generated as described before using CRISPR/CAS9 (31).

### Mouse strains

Experiments were performed in accordance with the Swiss federal regulations and approved by the local ethics committee, Basel-Stadt, Switzerland (Approval 3036 and 3099). All animals were bred in-house at the Department of Biomedicine facility (University of Basel, Switzerland) in pathogen-free, ventilated HEPA-filtered cages under stable housing conditions of 45-65% humidity, a temperature of 21-25°C, and a a gradual light–dark cycle with light from 7:00 am to 5 pm. Mice were provided with standard food and water without restriction (License: 1007-2H).

Siglec-E^loxP^ mice were generated in collaboration with Biocytogen Company and LysM-Cre mice were generated as described before (42). To study the role of Siglec-E KO on LysM-Cre expressing cells, Tm(Siglec-E x LysM-Cre) C57BL/6 were generated by crossing LysMCre mice with Siglec-E^loxP^ mice.

### Patient samples

Tumor and blood samples were collected at the University Hospital Basel and buffy coats from healthy donors were obtained from the Blood Bank (University Hospital Basel, Switzerland). Sample collection and use of corresponding clinical data were approved by the local ethics committee in Basel, Switzerland (Ethikkommission Nordwestschweiz, EKNZ, Basel Stadt. Switzerland) and written informed consent was obtained from all donors before sample collection.

### Cell culture

Cell lines and primary cells were cultured at 37°C and 5% CO_2_ and regularly checked for mycoplasma contamination. All cell lines except HEKT293T were maintained in Dulbecco’s Modified Eagle Medium (DMEM, Sigma), supplemented with 10% heat-inactivated fetal bovine serum (FBS, PAA Laboratories), 1x MEM non-essential amino acid solution (Sigma) and 1% penicillin/streptomycin (Sigma).

HEK293T, all primary cells and co-cultures were maintained in Roswell Park Memorial Institute Medium (RPMI, Sigma) supplemented with 10% heat-inactivated fetal bovine serum (FBS, PAA Laboratories), 1x MEM non-essential amino acid solution (Sigma), 1 mM sodium pyruvate (Sigma), 0.05 mM 2-mercaptoethanol (Gibco) and 1% penicillin/streptomycin (Sigma).

### Tumor digest, splenocyte and PBMC isolation

To obtain single cell suspension, human and mouse tumors were mechanically dissociated and subsequently enzymatically digested using accutase (PAA Laboratories), collagenase IV (Worthington), hyaluronidase (Sigma) and DNase type IV (Sigma) for 1 h at 37°C under constant agitation. Afterwards, samples were filtered using a 70 µM cell strainer and washed. Precision counting beads (BioLegend) were added to all mouse tumors to calculate the number of cells per gram of tumor.

Human peripheral blood mononuclear cells (PBMCs) were isolated from buffy coats by density gradient centrifugation using Hisopaque-1077 (Millipore) and SepMate PBMC isolation tubes (StemCell) according to the manufacturer’s protocol followed by red blood cell lysis using RBC lysis buffer (eBioscience) for 2 min at RT. Subsequently, cells were washed with PBS and ready for further analysis.

For splenocyte isolation, freshly harvested murine spleens were mechanically dissociated by filtering them through a 100 µM filter. After washing, red blood cells were lysed as described above.

For murine PBMC analysis, blood from the tail vein of mice was collected on day 14 of the experiment by tail vein puncture. After washing, red blood cells were lysed as described above and samples were used immediately for lectin staining.

Single cells suspensions were used immediately or frozen for later analysis in liquid nitrogen (in 90% FBS and 10% DMSO).

### Tumor models

Siglec-ExLysM-Cre mice (SigE^ΔLysM^) were injected subcutaneously into the right flank with 500 000 B16F10 melanoma, B16F10-sia, EL4 lymphoma or EL4 GFP cells in phenol red-free DMEM without additives. Siglec-E WT (SigE^WT^) sex-matched littermates were used as control. Mice were between 8-12 weeks of age at the beginning of the experiment and conditional knockout was confirmed by flow cytometry.

Tumor size was measured 3 times a week using a caliper. Animals were sacrificed before reaching a tumor volume of 1500 mm^3^ or when they reached an exclusion criterion. Tumor volume was calculated according to the following formula: Tumor volume (mm^3^) = (d^2^*D)/2 with D and d being the longest and shortest tumor parameter in mm, respectively.

### *In vivo* treatment

For *in vivo* Ly6G depletion, mice were injected intraperitoneally twice per week with 100 µg/mouse Ly6G depletion antibody (Clone: 1A8, BioXCell) in PBS. Injections were started one day before tumor injection and administered twice a week till mice reached the experimental endpoint.

For neutralization of CCL2 *in vivo*, mice were injected intraperitoneally 3 times a week with 200 µg/mouse CCL2 neutralization antibody (Clone: 2H5, BioXCell) in PBS. Antibody treatment was started one day after tumor injection and continued until the endpoint of the experiment.

### Multiparameter flow cytometry

Multicolor flow cytometry was performed on single cell suspension of cell lines, PBMCs, splenocytes or tumor digest. To avoid unspecific antibody binding, cells were blocked using rat anti-mouse FcγIII/II receptor (CD16/CD32) blocking antibodies (BD Bioscience) for murine and Fc Receptor Binding Inhibitor Polyclonal Antibody (Invitrogen) for human samples and subsequently stained with live/dead cell exclusion dye (Zombie Dyes, BioLegend). Surface staining was performed with fluorophore-conjugated antibodies (Table S1) or lectins for 30 minutes at 4°C in FACS buffer (PBS, 2% FCS, 0.5 mM EDTA). Stained samples were fixed using IC fixation buffer (eBioscience) until further analysis. For intracellular staining, cells were fixed and permeabilized using the Foxp3/transcription factor staining buffer set (eBioscience) and 1x Permeabilization buffer (eBioscience) according to the manufacturer’s instruction. All antibodies were titrated for optimal signal-to-noise ratio. Compensation was performed using AbC Total Antibody Compensation Bead Kit (Invitrogen) or cells. Samples were acquired on LSR II Fortessa flow cytometer (BD Biosciences), CytoFLEX (Beckmann Coulter) or Cytek Aurora (Cytek Biosciences) and analyzed using FlowJo 10.8 (TreeStar Inc). Cell sorting was performed using a BD FACSAria III or BD FACSMelody (BD Bioscience). Doublets, cell debris and dead cells were excluded before performing downstream analysis. Fluorescence-minus-one (FMO) samples were used to define the gating strategy and calculate mean fluorescence intensity (MFI).

To access desialylation status, cells were stained with lectins as described above. Fluorophore-coupled lectins - PNA-PE (GeneTex) and SNA-FITC (GeneTex) - and biotinylated lectins – MALII (GeneTex) were used at a final concentration of 10 µg/mL. Biotinylated lectins were detected using PE-Streptavidin (Biolegend).

### *In vitro* generation of human suppressive myeloid cells

To generate suppressive human myeloid cells, we used an adapted version of the protocol established by Lechner et al. (23).

#### A. Generation of MDSC-like cells

For *in vitro* MDSC induction, freshly isolated PBMCs from healthy donor buffy coats were co-cultured for 7 days with different cancer cell lines (A549, A549-GNE KO, A549-sia, HeLa, HeLa-sia) at a ratio of 1:100 in complete RPMI medium supplemented with 10 ng/mL GM-CSF (PeproTech). Cancer cells were seeded at an initial concentration of 1×10e4 cells/mL and the same amount of medium supplemented with GM-CSF was added at day 4 of the experiment. After one week, all confluent and adherent cells were collected using 0.05% trypsin-EDTA (Gibco).

#### D. Isolation of MDSC-like cells

For MDSC isolation, CD33 magnetic isolation was performed using the human CD33 positive selection kit II (StemCell) was used following the manufacturer’s instructions. Isolated cells were resuspended in complete RPMI medium used freshly for the suppression assay.

#### C. Isolation of autologous CD8 T cells

Autologous CD8 T cells were obtained from frozen PBMCs from the same donor using the CD8+ Microbeads human T cell isolation Kit (Miltenyi Biotec). To monitor cell proliferation, cells were labeled with 1.25 µM CellTraceViolet (Invitrogen) according to the manufacturer’s instructions. Washed cells were resuspended in complete RPMI and used for the suppression assay.

#### D. Suppression assay

Isolated MDSC-like cells and autologous CD8 T cells were co-cultured at indicated ratios for 5 days in a U-bottom plate in complete RPMI. T cells were stimulated by the addition of 100 IU/mL IL-2 (Proleukin) and anti-CD3/CD28 stimulation using loaded MACSiBead particles (Miltenyi Biotec) in a proportion of 1:1 of beads to cell. Unstimulated T cells and stimulated T cells without MDSC addition were used as controls. After five days, supernatants were frozen at -80°C, and the cells were stained for flow cytometry.

For Siglec-9 blocking, Siglec-9 blocking antibody (Clone 191240, R&D Systems) was added at a final concentration of 10 µg/mL. For sialidase treatment, MDSC were pretreated and washed before being added to the assay as described below.

### Human intratumoral-derived MDSC suppression assay

PBMCs and tumor digests were used freshly immediately after isolation as described above. MDSCs were isolated from tumor digest and CD8 T cells from PBMCs using CD33 Microbeads (Miltenyi) or CD8 Microbeads (Miltenyi), respectively. Cells were co-cultured at a MDSC:target ratio of 1:4 in a U-bottom plate for 5 days in complete RPMI in the presence of 30 IU IL-2 (Proleukin) and human CD2/CD3/CD28 T cell activator at a final concentration of 25 µL/mL (Immunocult, StemCell). For Siglec-9 blocking, Siglec-9 blocking antibody (Clone 191240, R&D Systems) was added at a final concentration of 10 µg/mL. For sialidase treatment, MDSC were pretreated and washed before being added to the assay as described below.

Supernatants were frozen at -80°C, and the cells were stained for flow cytometry.

### Murine MDSC suppression assay

Murine T cells were enriched from wildtype mouse splenocytes by negative selection using the murine pan T cell isolation Kit (EasySep, StemCell). To monitor T cell proliferation, isolated T cells were stained with 2.5 µM CellTraceViolet (Invitrogen) according to the manufacturer’s instructions.

Murine MDSCs were isolated from splenocytes of tumor-bearing mice by negative selection using a murine MDSCs isolation Kit (EasySep, StemCell). As indicated, obtained MDSCs were used immediately or pretreated with sialidase as described below.

Isolated MDSCs and T cells were plated in a ratio of 1:1 on a 96 well flat bottom plate and co-cultured for 48 hours in complete RPMI in the presence of 50 IU IL-2 (Proleukin). For T cell stimulation, the plate was coated with anti-CD3 (clone 17A2, BioLegend) and anti-CD28 (clone 37.51, BD Biosciences). Supernatants were frozen at -80°C, and the cells were stained for flow cytometry.

For Siglec-E blocking, purified anti-mouse Siglec-E antibody (M1305A02, BioLegend) was added at a final concentration of 10 µg/mL. CCL2 blocking was performed by the addition of 50 µg/mL CCL2 (Clone 2H5, BD).

### Sialidase treatment

To cleave terminal sialic acid residues, cells were treated with bacterial sialidase (*Vibrio cholerae*, Sigma) at a concentration of 10 µM for 20 minutes in PBS. Subsequently, cells were washed with complete medium and used for downstream analysis. Additionally, viral sialidase (active *H1N1*, Sino Biological) and bacterial sialidase (*Arthrobacter ureafaciens*, Roche) were used for pre-treatment which is indicated in the figure legends. If not stated otherwise, *Vibrio cholerae* bacterial sialidase was used.

### Lentivirus production and lentiviral transduction of cells lines

To generate cell lines expressing *H1N1* viral sialidase and GFP, A549, HeLa, B16F10 and EL4 were stably transduced with lentivirus.

For lentiviral production, 14×10e6 HEK293T cells were seeded 24 hours before transfection in 18 mL complete RPMI medium in a 15-cm culture dish. For the transfection mix, 1.9 µg pMD2.G, 3.5 µg pCMVR8.74 and 5.4 µg pLV transfer vector were mixed in 1.8 ml jetOPTIMUS buffer (Polyplus). 16.2 µl jetOPTIMUS (Polyplus) was added followed by 10 minutes of incubation and finally the addition of the prepared transfection mix. Medium was exchanged after 16 hours and lentiviral particles were collected 24 and 48 hours after medium exchange. The pooled supernatant was concentrated with 4x in-house made PEG-8000 solution and resuspended in PBS with 1% human serum albumin. Aliquots of the produced virus were stored at -80°C until further use. pMD2.G and pCMVR8.74 were kindly provided by Didier Trono (Addgene plasmid #12259 & #22036).

For lentiviral transduction, 50 000 cancer cells were seeded in a 24-well plate in 500 µL complete RPMI medium and rested overnight. Media was renewed with addition of 100 µL concentrated lentivirus and 8 µg/mL polybrene (Sigma). To increase transduction efficiency, spinoculation was performed and cells were centrifuged for 90 minutes at 800xg. Afterwards, cells were incubated at standard cell culture conditions and transduction efficiency was frequently checked by flow cytometry staining assessing sialidase expression, PNA, MALII and SNA levels.

### Cytokine and chemokine analysis

Collected supernatants from murine and human co-cultures were thawed on ice and aliquots were sent on dry ice to Eve Technologies (Canada). Cytokine and chemokine concentrations were analyzed and calculated by Eve Technology. For visualization, normalized values (z-scores) of each cytokine were calculated based on the mean and standard deviation of each marker.

### Bulk RNA Sequencing

MDSC-like cells generated with A549 and A549-sia cancer cells from 4 different healthy donors were harvested after 7 days of co-culture as described above. For purification of MDSCs, cells were stained with CD33-PE (Miltenyi) followed by CD33-positive selection using the EasySep human PE positive selection Kit II (StemCell). To increase purity, cells were further sorted for PE positivity by Aria III (BD Bioscience) flow cytometer. For RNA purification, sorted cells were washed and RNA isolated using the RNeasy Plus Micro Kit (Qiagen) including a gDNA elimination step by QIAshredder spin columns (Qiagen).

Quality control (QC length profiling and concentration using RiboGreen) and library preparation (TruSeq stranded mRNA HT Kit by Illumina) were performed by the Genomics Core Facility of the University Basel. Sequencing was performed on four lanes of the Illumina NextSeq 500 instrument resulting in 38nt-long paired-end reads. The dataset was analyzed by the Bioinformatics Core Facility, Department of Biomedicine, University of Basel. cDNA reads were aligned to ‘hg38’ genome using Ensembl 104 gene models with the STAR tool (v2.7.10a) with default parameter values except the following parameters: outFilterMultimapNmax=10, outSAMmultNmax=1, outSAMtype=BAM SortedByCoordinate, outSAMunmapped=Within. At least 40M read pairs were mapped per sample. The software R (v4.1.1) and the tool featureCounts from Subread (v2.0.1) package from Bioconductor (v3.14) were used to count aligned reads per gene with default parameters except: -O, -M, --read2pos=5, --primary, -s 2, -p, -B. Further analysis steps were performed using R (v4.2.0) and multiple packages from Bioconductor (v3.15). The package edgeR (v3.38.1) was used to perform differential expression analysis. A gene was included in the analysis only if it had at least 1 count per million (CPM) in at least any four samples. Gene set enrichment analysis was performed using the tool camera from the edgeR package and the Gene Ontology gene set (category C5) from MSigDB (v7.5.1).

### N-Glycomics analysis

For N-glycomic profiling, harvested cells were extracted with lysis buffer containing 7 M urea, 2 M thiourea, 10 mM dithioerythreitol in 40 mM Tris buffer with 1% protease inhibitor (Roche). The cell membranes were disrupted by High Intensity Focused Ultrasound with 10 times 10 s sonication with 16 amplitudes and 1 minute on ice in between, and subsequent shaking for 4 hours at cold room. The protein extracts were alkylated with 100 mM iodoacetamide in the dark for 4 hours at 37°C. Ice-cold trichloroacetic acid was added to a final 10% w/v concentration and left for one hour. After centrifugation at 20000 g for 30 minutes at 4°C, precipitated sample pellets were washed twice with ice-cold acetone and then lyophilized. Dry protein pellets were redissolved in 50 mM ammonium bicarbonate buffer (pH 8.5), 250 unit of benzonase nuclease (Sigma-Aldrich) was added and incubated for 30 minutes at 37°C, following by trypsin digestion overnight. After deactivating the activity of trypsin, protein mixtures were further treated with PNGaseF (New England Biolab). The released glycans were cleaned up according to previous studies (43).

For MALDI-MS analyses, the glycan samples were permethylated using the sodium hydroxide/dimethyl sulfoxide slurry method, as described by Dell et al (44). The samples were dissolved in 20 µL of acetonitrile. 1 µL sample mixed with 10 mg/mL 2,5-Dihydroxybenzoic acid (Bruker) in 70% Acetonitrile with 1mM sodium chloride was spotted on MALDI target plate and analyzed by Bruker RapiFlex^TM^ MALDI-TOF-TOF. Permethylated high mannose N-glycans and glycans from fetuin were used to calibrate the instrument prior the measurement. The laser energy for each analysis was fixed and the data were accumulated from 10 000 shots. The data was analyzed by GlycoWorkbench (45) and inspected manually. For relative quantification, the data was first deisotoped and the peak height was used for the calculation based on the following equation:

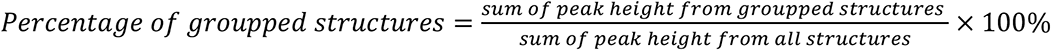

### Statistical Analysis

All statistical analysis were performed using GraphPad Prism9. Used statistical tests as well as sample sizes are indicated in the figure legend.

p values > 0.05 were considered not significant, p values < 0.05 were considered significant. Asterisks indicate: * p value < 0.05, ** p value < 0.01, *** p value < 0.001, **** p value < 0.0001. n indicates the number of biological replicates, all bars within the graphs represent mean values, and the error bars represent standard errors of the mean (SEM) or standard deviation (SD) as indicated.

## Supplementary Figures

**Supp. Figure 1:**
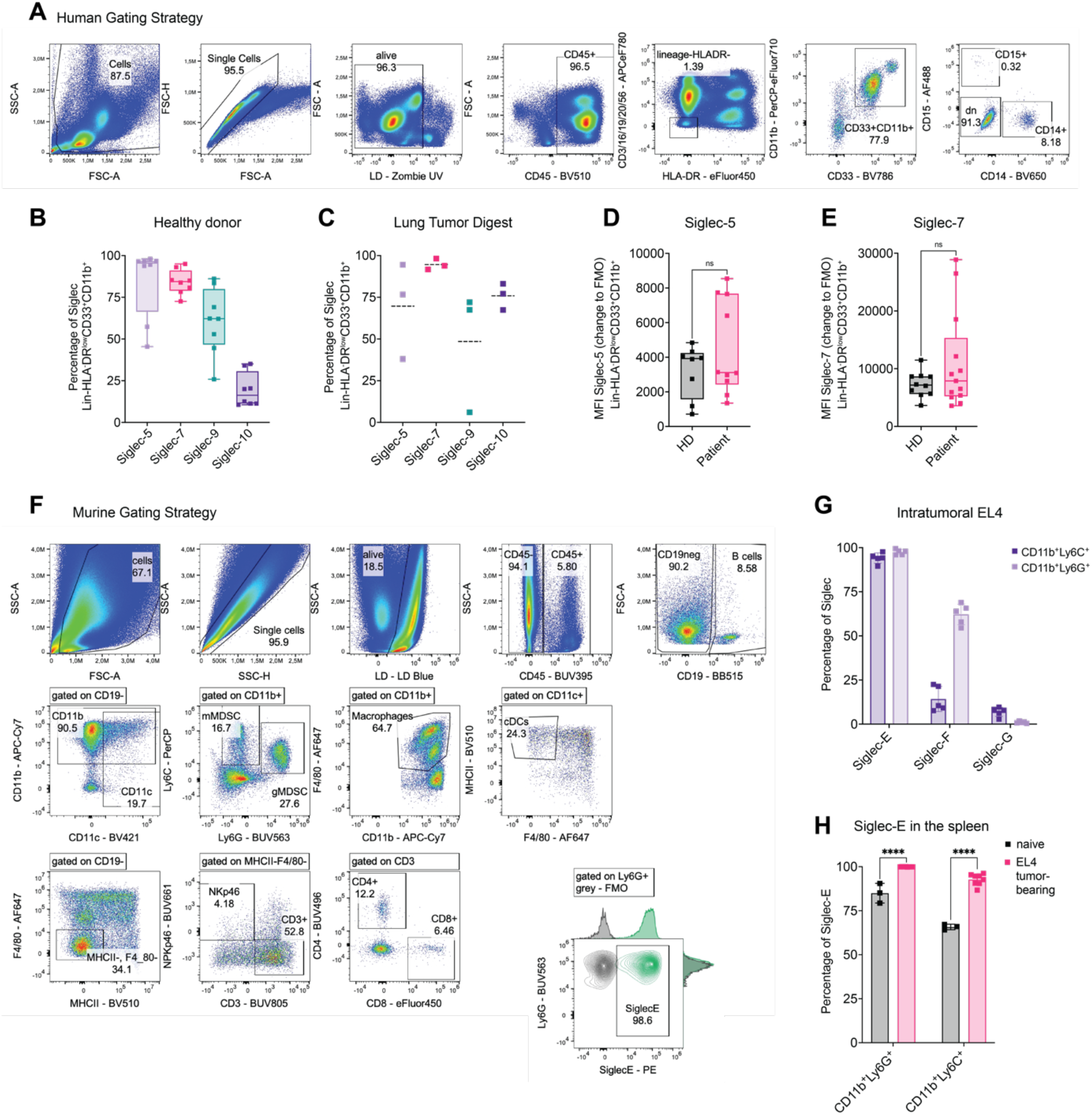
Expression of Siglecs on myeloid cells from various origins. **(A)** Exemplary gating strategy used for Fig 1A-E. MDSCs were gated as CD45^+^Lin^-^(CD3^-^CD16^-^ CD19^-^CD20^-^CD56^-^) HLADR^low^CD33^+^CD11b^+^ cells. **(B)** Percentage of Siglec-5, Siglec-7, Siglec-9 and Siglec-10 expressed on CD45^+^Lin^-^HLADR^low^CD33^+^CD11b^+^ cells derived from healthy donor peripheral blood (PB) or **(C)** intratumorally from lung cancer tumor digest. *N=3-8 donors with at least N=2* **(D)** MFI of Siglec-5, **(E)** Siglec-7 gated on CD45^+^Lin^-^HLADR^low^CD33^+^CD11b^+^ cells derived from healthy donor and lung cancer patient PB. MFI is shown as change to FMO and was determined by flow cytometry. *N=8-13 donors with at least N=2 experiments.* **(F)** Representative gating strategy to identify murine immune cell types in tumor digest, peripheral blood and spleen. **(G)** Subcutaneously injected endpoint tumors from EL4 lymphoma engrafted mice were harvested, digested and immune cell infiltration was assessed by multi-color flow cytometry. Siglec-E, Siglec-F and Siglec-G expression was assessed on CD45^+^CD11b^+^Ly6G^+^ and CD45^+^CD11b^+^Ly6C cells. *N=5 mice.* **(H)** Spleens from naïve and EL4 lymphoma tumor-bearing mice at endpoint were collected and analyzed for Siglec-E expression via flow cytometry. *N=3-8 mice per group*. Data are presented as mean. Error bar values represent SD. Two-tailed unpaired Student’s t test or multiple unpaired t-tests (H) was used. *P<0.05, **P<0.01, ***P<0.001, and ****P<0.0001.

**Supp. Figure 2:**
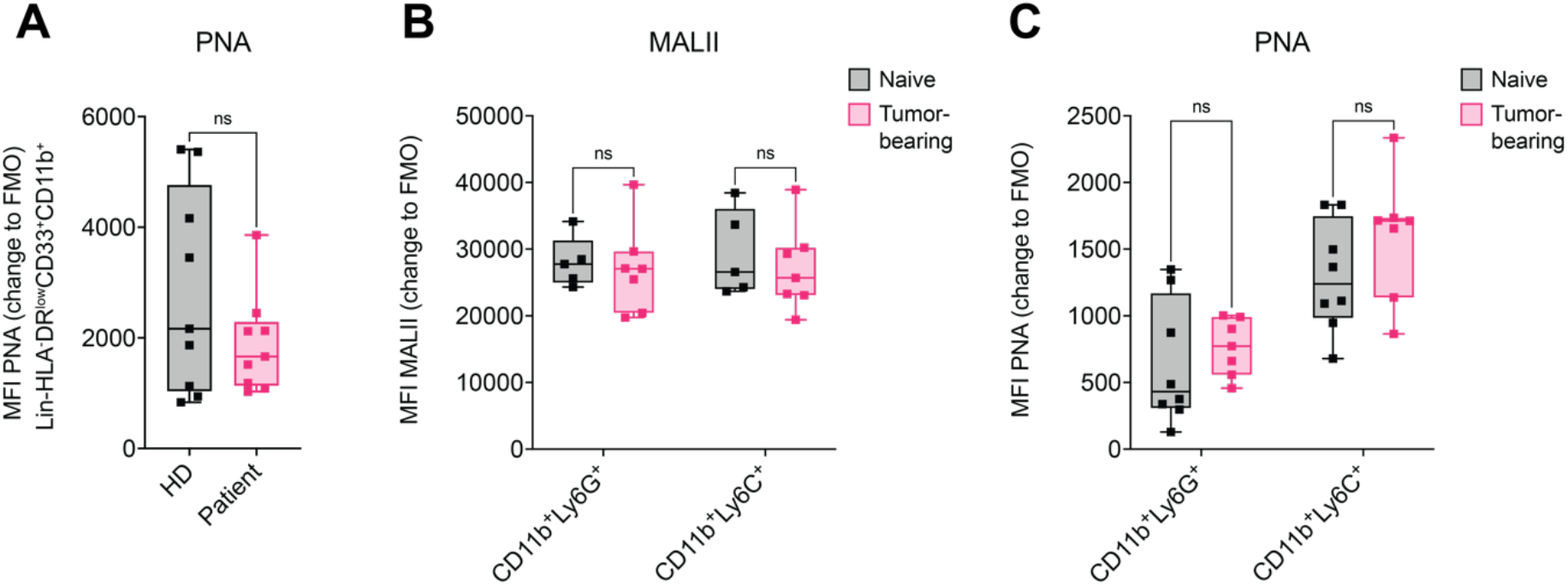
Sialoglycan expression on myeloid cells in human and mice. **(A)** PNA gated on PB-derived CD45^+^Lin^-^HLADR^low^CD33^+^CD11b^+^ cells from primary lung cancer patient and healthy controls. MFI is shown as a change to FMO and was determined by flow. *N=8-12 donors with at least N=2*. **(B)** Fresh blood from B16F10 tumor-bearing mice and naïve wildtype mice was collected at day 14 after tumor inoculation and analyzed for MALII or **(C)** PNA gated on CD45^+^CD11b^+^Ly6C^+^ or CD45^+^CD11b^+^Ly6G^+^ cells. MFI is shown as a change to FMO. *5-8 mice per group.* Data are presented as mean. Error bar values represent SD. Two-tailed unpaired Student’s t test or multiple unpaired t-tests (B, C) was used. *P<0.05, **P<0.01, ***P<0.001, and ****P<0.0001.

**Supp. Figure 3:**
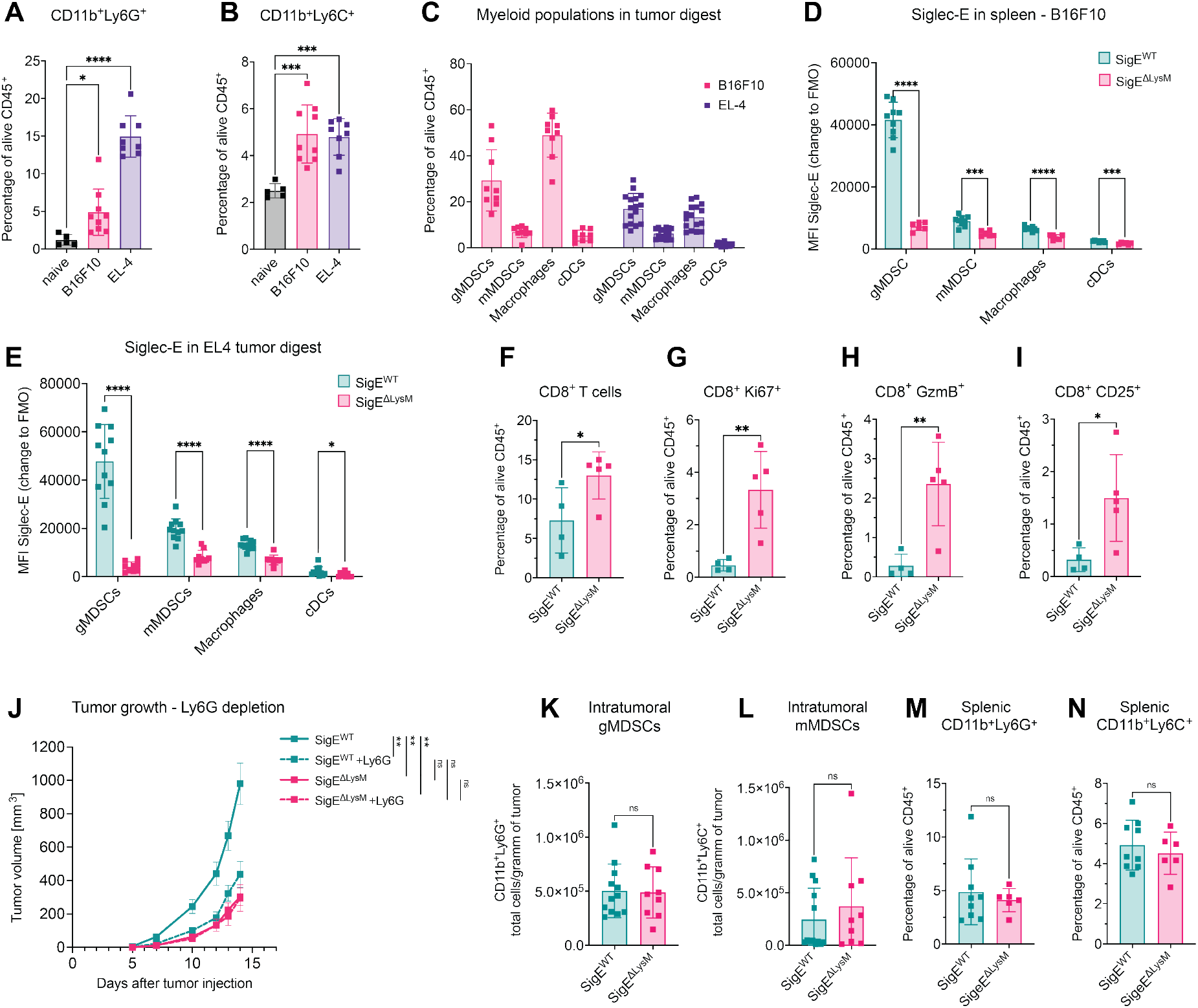
Siglec-E depletion on myeloid cells leads to survival benefits in tumor models of emergency myelopoiesis. **(A)** Spleens from naïve mice, EL4 lymphoma or B16F10 melanoma tumor-bearing mice at endpoint were collected and analyzed for CD11b^+^Ly6G^+^ and **(B)** CD11b^+^Ly6C^+^ cell infiltration. *N=5-9 mice per group*. **(C)** Tumor digest analyzed at endpoint for myeloid cell infiltration. gMDSCs (CD45^+^CD11b^+^Ly6G^+^), mMDSCs (CD45^+^CD11b^+^Ly6C^+^), macrophages (CD45^+^CD11b^+^F4/80^+^), dendritic cells (DCs) (CD45^+^CD11c^+^MHCII^+^F4/80^-^) are shown as percentage of CD45 cells. *N=9-16 mice per group*. **(D)** MFI of Siglec-E expression in B16F10-tumor bearing mice was analyzed on myeloid cells in the spleen at endpoint of the experiment. *N=6-9 mice per group*. **(E)** MFI of Siglec-E expression in EL4 tumor-bearing mice was analyzed on myeloid cells in tumor digest at endpoint*. N=4-5 mice per group*. **(F)** Subcutaneous EL4-GFP tumors were analyzed at endpoint via flow cytometry. Intratumoral CD8 T cells at endpoint (CD45^+^aliveCD19^-^NKp46^-^CD3^+^CD8^+^) were further sub gated on **(G)** Ki67^+^, **(H)** GranzymeB^+^(GzmB^+^) **(I)** CD25^+^ CD8 T cells and quantified as percentage of CD45^+^ cells*. N=4-5 mice per group*. **(J)** Tumor volume from pooled data from 2 independent experiments. Siglec-E^ΔLysM^ mice and Siglec-E^WT^ mice were treated up to 6 times with Ly6G depletion antibody as shown in Figure 2 I. *N=9-11 mice per group.* **(K)** Tumor digest analyzed at endpoint for myeloid cell infiltration. gMDSCs (CD45^+^CD11b^+^Ly6G^+^) and **(L)** mMDSCs (CD45^+^CD11b^+^Ly6C^+^) were quantified as cells per gram of tumor at the endpoint of the experiment. Pooled data from 2 independent experiments. *N=9-12 mice per group.* **(M)** Spleens from B16F10 melanoma tumor-bearing mice at endpoint were collected and analyzed for CD11b^+^Ly6G^+^ and **(N)** CD11b^+^Ly6C^+^ cells. Cells were quantified as percentage of alive CD45^+^ cells. *N=6-9 mice per group.* Data are presented as mean. Error bar values represent SD or SEM (F, G, L). Two-tailed unpaired Student’s t test, multiple unpaired t-tests (D, E) or one-way ANOVA followed by Dunnett’s multiple comparisons test (A, B) was used. Tumor growth was compared by mixed-effects analysis followed by Bonferroni’s multiple comparisons test. *P<0.05, **P<0.01, ***P<0.001, and ****P<0.0001.

**Supp. Figure 4:**
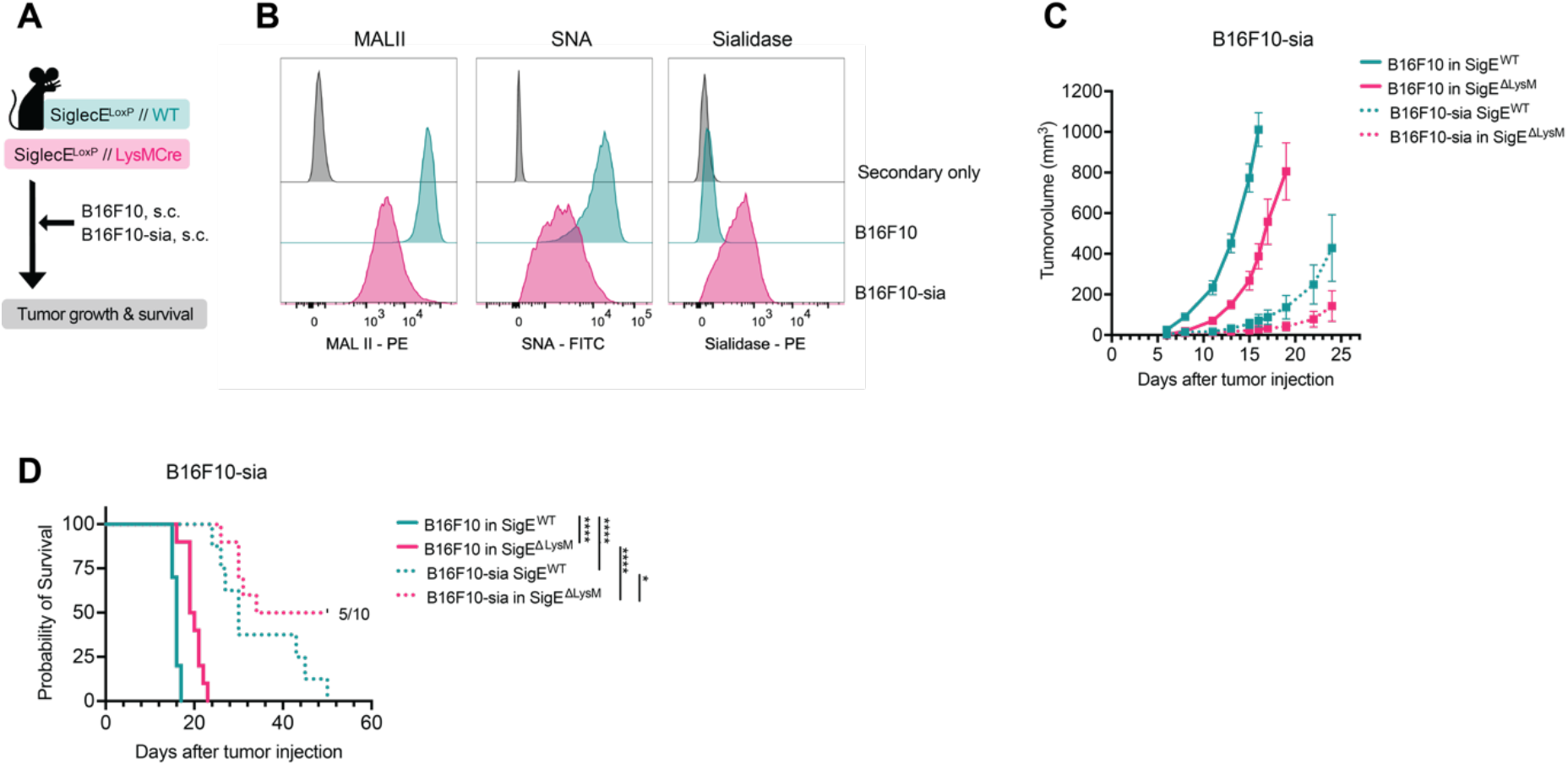
Combination of sialidase expression and lack of Siglec-E on myeloid cells prolongs survival *in vivo*. **(A)** Experimental setup: SigE^ΔLysM^ and SigE^WT^ littermates were subcutaneously injected with B16F10 or B16F10 cells expressing sialidase (B16F10-sia). Tumor growth and probability of survival were addressed as the main read-out. **(B)** B16F10 and B16F10-sia cells were stained for SNA, MALII and Sialidase expression to validate the successful generation of stable cell lines. Cell lines were stained before each experiment, representative results are shown. **(C)** Tumor volume and **(D)** Kaplan-Meier survival curves from pooled data from 2 independent experiments. *N=8-12 mice per group from 2 experiments.* Data are presented as mean with error bars presenting SEM. Tumor growth was compared by mixed-effects analysis followed by Bonferroni’s multiple comparisons test. For survival analysis, log-rank test was used followed by Šidák correction for multiple comparisons. *P<0.05, **P<0.01, ***P<0.001, and ****P<0.0001.

**Supp Figure 5:**
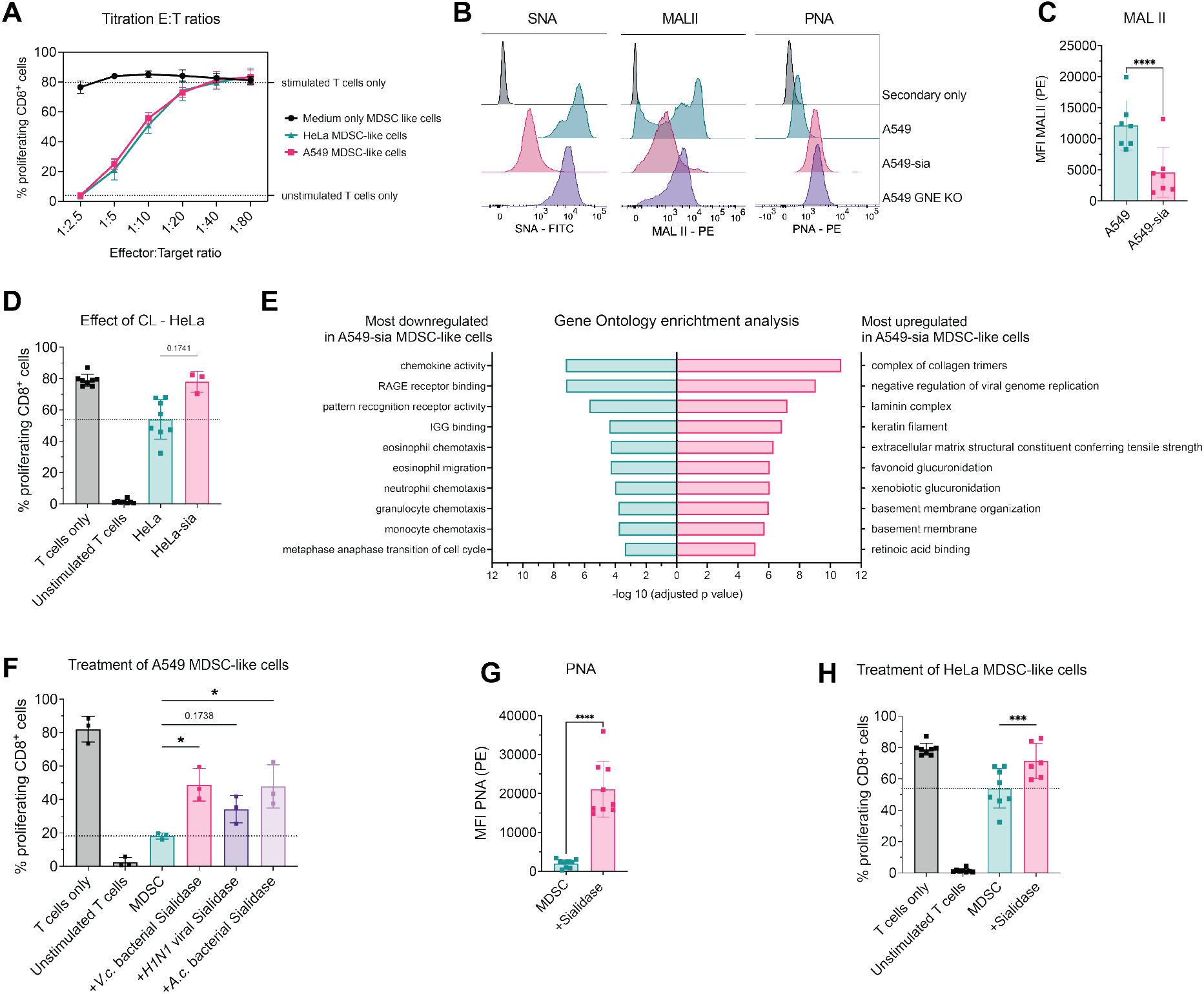
Sialylation modulates the suppressive potential of human CD33^+^ cells. **(A)** Identification of E:T ratios to validate the suppressive capacity of CD33^+^ cells against CD8^+^ cells. Suppressive myeloid cells were generated by co-culture with A549 (pink), HeLa (green) or without cancer cells (black). Dotted lines indicate the proliferation of T cells alone with/without stimulation by IL-2, aCD3/28 microbeads. *N=4 donors of N=2 experiments*. **(B)** A549, A549 expressing sialidase (A549-sia) and A549 GNE KO cells were stained for SNA, MALII and PNA to validate the successful generation of stable cell lines. Cell lines were stained before each experiment, representative results are shown. **(C)** MALII staining was performed on suppressive CD33^+^ cells on day 7 of the experiment. N=7 donors of *N=4 experiments.* **(D)** Percentage of proliferating CD8^+^ cells upon co-culture (1:10 ratio) with indicated suppressive CD33^+^ cells. Suppressive myeloid cells were generated using HeLa or HeLa-expressing sialidase (HeLa-sia) cancer cell lines. *N=3-8 donors* **(E)** Gene ontology enrichment analysis of the top 10 up- and downregulated gene sets found in suppressive CD33^+^ cells generated with A549-sia compared to parental A549 cell line. **(F)** PNA staining was assessed on suppressive CD33^+^ cells after pretreatment with sialidase on day 7 of the experiment. *N=9 donors of N=5 experiments.* **(G)** Proliferating CD8^+^ cells in percentage co-cultured with suppressive CD33^+^ cells generated by A549 co-culture in a ratio of 1:5. CD33^+^ cells were used immediately or were pretreated with indicated sialidases. *N=3 donors of N=2 experiments.* **(H)** Percentage of proliferating CD8^+^ cells upon co-culture (1:10 ratio) with suppressive CD33^+^ cells generated by HeLa co-culture. CD33^+^ cells were used immediately or were pretreated with sialidase. Data are presented as mean and error bar values represent SD. Paired t-test was used. *P<0.05, **P<0.01, ***P<0.001, and ****P<0.0001.

**Supp. Figure 6:**
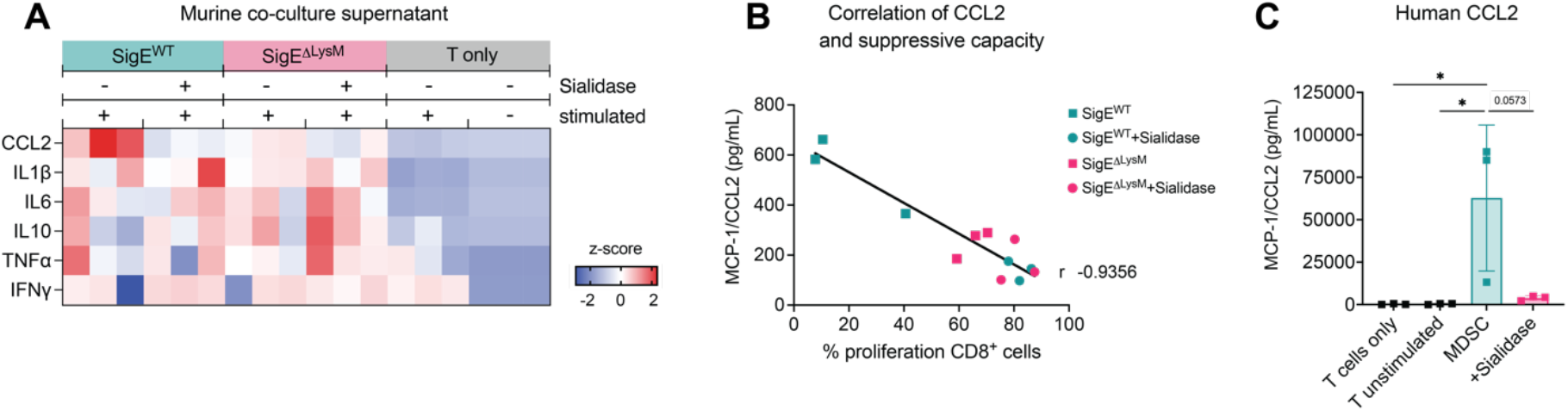
Suppressive myeloid cells correlate with CCL2 expression in humans and mice. **(A)** Cytokines found in the supernatant of murine MDSC:T cell co-cultures at endpoint of the experiment from figure 3. Z-scores were calculated for each cytokine. MDSCs were untreated, pretreated with sialidase or Siglec-E was added to the co-culture. *N=3 donors per group*. **(B)** Correlation of MCP-1 levels measured in supernatants of murine MDSC:T cell co-cultures at endpoint and percentage of proliferation of CD8^+^ T cells from the same condition. *N=3 donors per group*. **(C)** MCP-1/CCL2 found in the supernatant of human primary CD33^+^:CD8^+^ cell co-cultures at endpoint of the experiment from figure 4H. CD33^+^ cells were untreated or pretreated with sialidase. Z-scores were calculated for each cytokine and are shown on a color scale from blue to red. *N=3 donors per group*. Data are presented as mean and error bar values represent SD. One-way ANOVA was used. R shows the Pearson correlation coefficient. *P<0.05, **P<0.01, ***P<0.001, and ****P<0.0001.

**Table S1:**
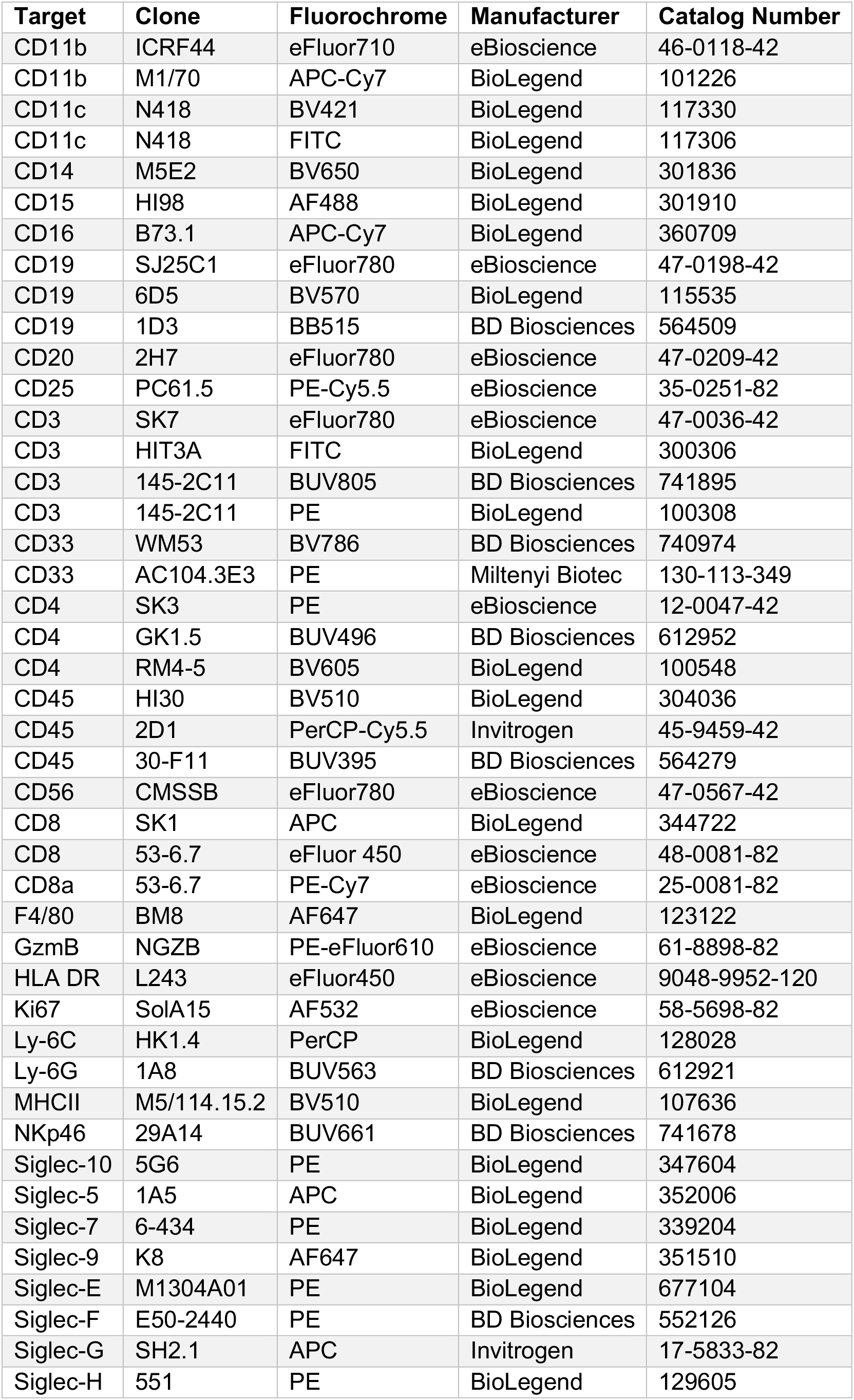
Flow cytometry antibodies.

| **Target** | **Clone** | **Fluorochrome** | **Manufacturer** | **Catalog Number** |
| --- | --- | --- | --- | --- |
| CD11b | ICRF44 | eFluor710 | eBioscience | 46-0118-42 |
| CD11b | M1/70 | APC-Cy7 | BioLegend | 101226 |
| CD11c | N418 | BV421 | BioLegend | 117330 |
| CD11c | N418 | FITC | BioLegend | 117306 |
| CD14 | M5E2 | BV650 | BioLegend | 301836 |
| CD15 | HI98 | AF488 | BioLegend | 301910 |
| CD16 | B73.1 | APC-Cy7 | BioLegend | 360709 |
| CD19 | SJ25C1 | eFluor780 | eBioscience | 47-0198-42 |
| CD19 | 6D5 | BV570 | BioLegend | 115535 |
| CD19 | 1D3 | BB515 | BD Biosciences | 564509 |
| CD20 | 2H7 | eFluor780 | eBioscience | 47-0209-42 |
| CD25 | PC61.5 | PE-Cy5.5 | eBioscience | 35-0251-82 |
| CD3 | SK7 | eFluor780 | eBioscience | 47-0036-42 |
| CD3 | HIT3A | FITC | BioLegend | 300306 |
| CD3 | 145-2C11 | BUV805 | BD Biosciences | 741895 |
| CD3 | 145-2C11 | PE | BioLegend | 100308 |
| CD33 | WM53 | BV786 | BD Biosciences | 740974 |
| CD33 | AC104.3E3 | PE | Miltenyi Biotec | 130-113-349 |
| CD4 | SK3 | PE | eBioscience | 12-0047-42 |
| CD4 | GK1.5 | BUV496 | BD Biosciences | 612952 |
| CD4 | RM4-5 | BV605 | BioLegend | 100548 |
| CD45 | HI30 | BV510 | BioLegend | 304036 |
| CD45 | 2D1 | PerCP-Cy5.5 | Invitrogen | 45-9459-42 |
| CD45 | 30-F11 | BUV395 | BD Biosciences | 564279 |
| CD56 | CMSSB | eFluor780 | eBioscience | 47-0567-42 |
| CD8 | SK1 | APC | BioLegend | 344722 |
| CD8 | 53-6.7 | eFluor 450 | eBioscience | 48-0081-82 |
| CD8a | 53-6.7 | PE-Cy7 | eBioscience | 25-0081-82 |
| F4/80 | BM8 | AF647 | BioLegend | 123122 |
| GzmB | NGZB | PE-eFluor610 | eBioscience | 61-8898-82 |
| HLA DR | L243 | eFluor450 | eBioscience | 9048-9952-120 |
| Ki67 | SolA15 | AF532 | eBioscience | 58-5698-82 |
| Ly-6C | HK1.4 | PerCP | BioLegend | 128028 |
| Ly-6G | 1A8 | BUV563 | BD Biosciences | 612921 |
| MHCII | M5/114.15.2 | BV510 | BioLegend | 107636 |
| NKp46 | 29A14 | BUV661 | BD Biosciences | 741678 |
| Siglec-10 | 5G6 | PE | BioLegend | 347604 |
| Siglec-5 | 1A5 | APC | BioLegend | 352006 |
| Siglec-7 | 6-434 | PE | BioLegend | 339204 |
| Siglec-9 | K8 | AF647 | BioLegend | 351510 |
| Siglec-E | M1304A01 | PE | BioLegend | 677104 |
| Siglec-F | E50-2440 | PE | BD Biosciences | 552126 |
| Siglec-G | SH2.1 | APC | Invitrogen | 17-5833-82 |
| Siglec-H | 551 | PE | BioLegend | 129605 |

## Abbreviations

A549-sia: A549 cells stably expressing viral *H1N1* sialidase
B16F10-sia: B16F10 cells stably expressing viral *H1N1* sialidase
DCs: Dendritic cells
DMEM: Dulbecco’s Modified Eagle Medium
E:T: Effector:target
gMDSC: Granulocytic myeloid-derived suppressor cell
GNE: UDP-N-acetylglucosamine 2-epimerase/N-acetylmannosamine kinase
GzmB: GranzymeB
HeLa-sia: HeLa cells stably expressing viral *H1N1* sialidase
ITIM: Immunoreceptor tyrosine-based inhibitory motifs
KO: Knockout
MALII: Maackia Amurensis Lectin II
MDSC: Myeloid-derived suppressor cell
MFI: Mean fluorescence intensity
mMDSC: Monocytic myeloid-derived suppressor cell
MS: Mass spectrometry
NK cells: Natural Killer cells
PBMCs: Peripheral blood mononuclear cells
PMN-MDSC: Polymorphonuclear myeloid-derived suppressor cell
PNA: Peanut Agglutinin
RPMI: Roswell Park Memorial Institute Medium
Siglec: Sialic acid-binding immunoglobulin-like lectin
SigE^ΔLysM^: Siglec-ExLysM-Cre; mice with knockout of Siglec-E on LysM expressing cells
SigE^WT^: Siglec-E wildtype
SNA: Sambucus Nigra Lectin
TAM: Tumor-associated macrophages
TME: Tumor microenvironment
V.c.: Vibrio cholerae

